# Methyl-CpG binding domain 2 (Mbd2) is an Epigenetic Regulator of Autism-Risk Genes and Cognition

**DOI:** 10.1101/247197

**Authors:** Elad Lax, Sonia DoCarmo, Yehoshua Enuka, Daniel M. Sapozhnikov, Lindsay A. Welikovitch, Niaz Mahmood, Shafaat A. Rabbani, Liqing Wang, Jonathan P. Britt, Wayne W. Hancock, Yosef Yarden, Moshe Szyf

**Author notes:** Corresponding author Moshe Szyf, Department of Pharmacology and Therapeutics, McGill University, 3655 Sir William Osler Promenade, Montreal, QC, Canada.

## Abstract

The Methyl-CpG-Binding Domain Protein family has been implicated in neurodevelopmental disorders. The Methyl-CpG-binding domain 2 (Mbd2) binds methylated DNA and was shown to play an important role in cancer and immunity. Some evidence linked this protein to neurodevelopment. However, its exact role in neurodevelopment and brain function is mostly unknown.

Here we show that *Mbd2*-deficiency in mice (*Mbd2*−/−) results in deficits in cognitive, social and emotional functions. Mbd2 binds regulatory DNA regions of neuronal genes in the hippocampus and loss of *Mbd2* alters the expression of hundreds of genes with a robust down-regulation of neuronal gene pathways. Further, a genome-wide DNA methylation analysis found an altered DNA methylation pattern in regulatory DNA regions of neuronal genes in *Mbd2*−/− mice. Differentially expressed genes significantly overlap with gene-expression changes observed in brains of Autism Spectrum Disorder (ASD) individuals. Notably, down-regulated genes are significantly enriched for human ortholog ASD risk-genes. Observed hippocampal morphological abnormalities were similar to those found in individuals with ASD and ASD rodent models. Hippocampal *Mbd2* knockdown partially recapitulates the behavioral phenotypes observed in *Mbd2*−/− mice.

These findings suggest Mbd2 is a novel epigenetic regulator of genes that are associated with ASD in humans. Mbd2 loss causes behavioral alterations that resemble those found in ASD individuals.

## Introduction

Epigenetic modifications of the genome are long known to play a crucial role in normal brain function including a wide range of neuropsychological processes as well as in neuropsychological disorders [1]. The most studied epigenetic modification is DNA methylation, the addition of a methyl group to the DNA on a cytosine in a CpG dinucleotide context. DNA methylation in promoters and other regulatory regions suppress gene expression by interfering with transcription-factors and transcription machinery binding [2, 3]. An additional mechanism involves recruitment of members of a methylated-DNA binding proteins family (MECP2 and MBD1-6) which share a Methyl-CpG Binding Domain (MBD) [4, 5]. MECP2 and MBD2 were shown to recruit chromatin repressive complexes to genes and thus cause alterations in histone modification and silencing of gene-expression [6–8]. These proteins are highly expressed in brain tissues [9, 10].

The most extensively studied MBD protein is MeCP2 since mutations and duplications of this gene cause Rett syndrome [11, 12]. Some studies on the role of MBD1 suggest it has a role in neurodevelopment and neurodevelopmental disorders like autism [13]. In contrast, little is known about the roles of other MBD proteins in the brain.

Several studies associated MBD2 with neuropsychiatric disorders. An increased MBD2 DNA binding on the promoters of *BDNF, GAD1* and *RELN* genes was observed in post-mortem brains of schizophrenia and bipolar disorder patients [14]. There is evidence for rare nonsynonymous de novo mutations in *MBD2* identified in ASD [15–17]. Several studies found Copy Number Variants (CNV) at and around the genomic position of the MBD2 gene (18q21.2) in ASD individuals (both deletions and duplications; see SFARI-CNV database: https://gene.sfari.org/database/cnv/). However, in these cases the CNVs span also other genes associated with ASD and related disorders, including *TCF4* (Pitt-Hopkins syndrome), ASXL3 (Bainbridge-Ropers syndrome) and *SETBP1* (ID syndrome MIM #616078 and Schinzel-Giedion syndrome); thus confounding the role of Mbd2 deletions in the etiology of ASD.

MBD2-bound DNA regions are hypo-methylated in the prefrontal cortex of depressed suicide-completers [18]. In rat pups of low maternal care there is reduced Mbd2 expression in the hippocampus which corelates with reduced glucocorticoid receptor expression and elevated stress [19]. However, to date the mechanisms by which Mbd2 affect gene-expression that ultimately affect brain function and behaviors are unclear.

In the present study, we directly assessed the role of *Mbd2* in behavior using a knockout mouse model (*Mbd2*−/−). A comprehensive behavioral battery found cognitive, social and emotional deficits in Mbd2−/− mice. Several lines of evidence link ASD-associated MBD proteins to hippocampus development and function, including MeCP2 [6–22] and Mbd1 [13, 23]. Since the behavioral abnormalities pointed to hippocampal functions, we examined the molecular footprints of *Mbd2* in the hippocampus. We applied unbiased genome-wide approaches. Using ChIP-seq we mapped the genome binding sites of Mbd2 and found that it binds methylated and unmethylated CpGs on and around many neuronal genes in the hippocampus as well as other genomic loci. Loss of Mbd2 binding in *Mbd2*−/− mice led to down-regulation of neuronal-gene expression. In contrast, up-regulated genes in *Mbd2*−/− mice were associated mostly to homeostasis and cell maintenance gene pathways. *Mbd2*-deficiency led to increased methylation on promoters and enhancers of neuronal genes, suggesting that Mbd2 affects DNA methylation status at neuronal genes and hence hippocampal genome functions. Furthermore, we found that differentially expressed genes were highly overlapped and correlated to differentially expressed genes found in ASD individuals’ brain. Down-regulated genes were highly enriched for ortholog ASD risk-genes implying an important role for Mbd2 in neurodevelopment and neuropsychiatric disorders.

## Results

### Mbd2 is required for a range of cognitive, social and emotional behaviors

Previous studies on *Mbd2*−/− mice behavior found impaired maternal behaviors such as suboptimal feeding and delayed pup retrieval [24]. Hypoactivity, impaired nest-building behavior and mild spatial learning and memory impairments were found in male Mbd2−/− mice [25]. We further tested whether *Mbd2* is involved in behavior using additional behavioral tests. We tested both male and female mice and found no significant effect of sex for any of the behaviors measured (two-way ANOVAs, p>0.05 for main effect of sex and for interactions in all cases, FigS1–FigS3, with the exception of a significant main effect for sex on average grooming bout length (Fig S2C)). We therefore, grouped both males and females for subsequent experiments. There was no change in general locomotion in an open-field box (Fig 1A) or in other exploratory behaviors (speed, entries to the center of arena; Fig S4). However, we observed a significantly increased self-grooming time and number of grooming bouts in Mbd2−/− mice (Fig 1B-D). Several forms of memory were tested as well. Short-term working-memory was assessed by spontaneous alternations in Y-Maze and was found to be intact in *Mbd2*−/− mice. Also, the number of arm entries did not differ between groups suggesting normal exploration behavior in *Mbd2*−/− mice (Fig1E). In contrast, *Mbd2*−/− mice showed impaired memory-retention in the long-term object-location memory test compared to wild-type littermates and failed to explore the object in the novel location for more than chance levels (Fig1F). A previous study [25] did not find significant deficits in social behavior in *Mbd2*−/− mice using a three-chamber test which is a common test for sociability in mice. However, some mouse models for neurodevelopmental disorders show normal or near normal behavior in the three-chamber test while showing reduced sociability in the social interaction test [6–28] suggesting that this test is more sensitive. We therefore used the social-interaction test and found that *Mbd2*−/− mice exhibited reduced social-interaction time (Fig 1G). We also measured anxiety-like behavior in the dark-light box and found that *Mbd2*−/− mice spent less time in the light compartment which indicates higher anxiety levels (Fig1H). Our findings support the hypothesis that Mbd2 is involved in regulating cognitive, social and emotional behaviors.

**Figure 1.**
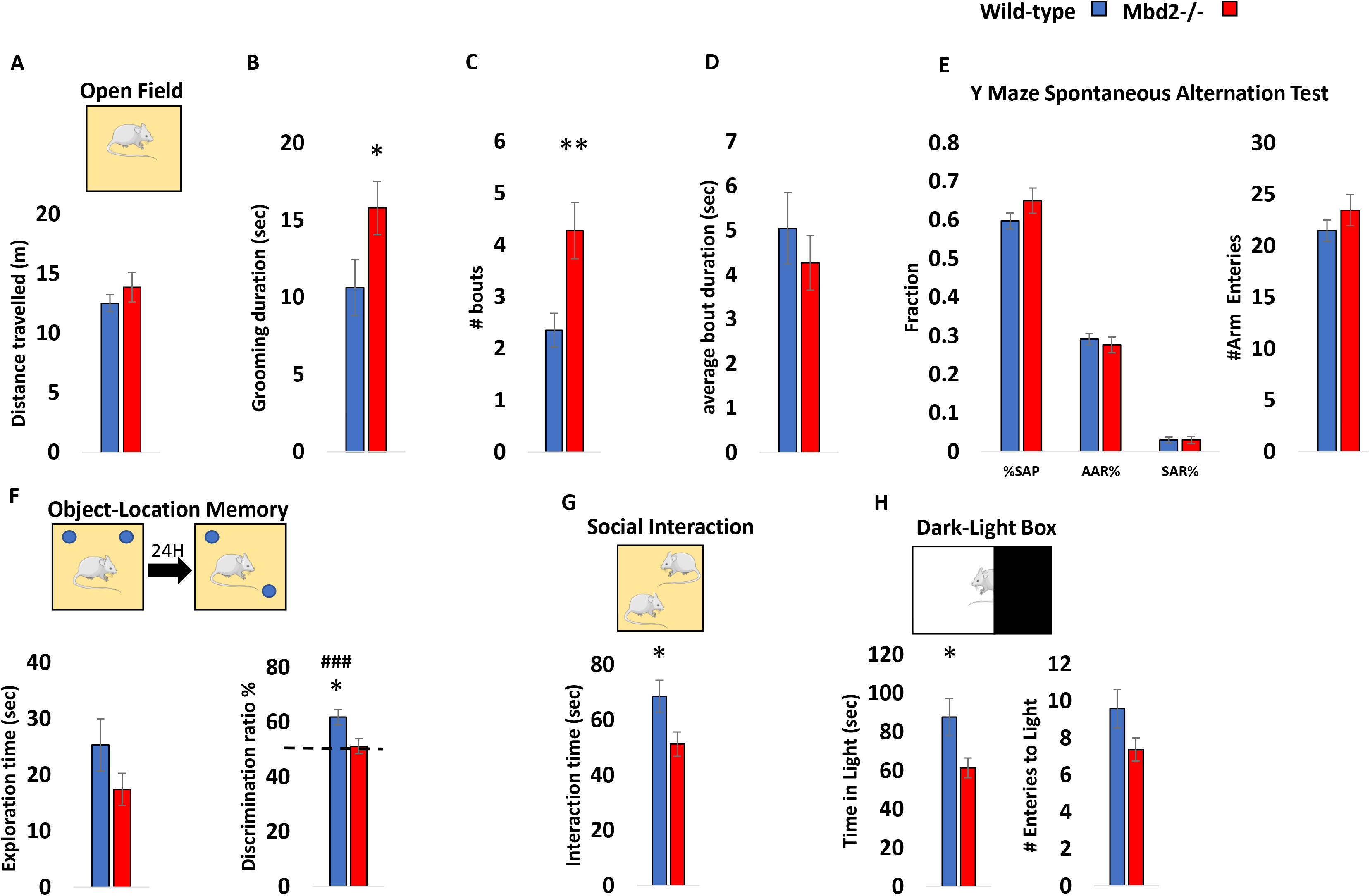
Behavioral effects of *Mbd2* deficiency. A. Locomotion in the Open-field box was assessed for 5 min. n=14 wild-type, n=15 mMbd2−/−. B. Self-grooming during open field. (Two-way ANOVA, main effect of genotype F(1,28)=5.2, p= 0.0304). C. Number of self-grooming bouts. (Two-way ANOVA, main effect of genotype F(1,28)=9.62, p= 0.0044). D. Average self-grooming bout duration. E. Y-maze spontaneous alteration was not affected in *Mbd2*−/− mice (right). SAP-Spontaneous Alteration Performance, AAR-Alternate Arm Return, SAR-Same Arm Return. Exploration is expressed as number of arm entries (left). n=17 wild-type, n=17 *Mbd2*−/−. F. Exploration time during object-location memory training (left) and discrimination ratio in object-location memory test (right). Discrimination ratio was significantly lower in *Mbd2*−/− mice (Two-way ANOVA, main effect of genotype F(1,24)=5.05, p= 0.0341; n=13 wild-type, n=15 *Mbd2*−/−). G. Social interaction. Mice were introduced to a novel mouse for 5 min and interaction time was recorded. (Two-way ANOVA, main effect of genotype F(1,29)=5.02, p= 0.0329; n=17 wild-type, n=16 *Mbd2*−/−). H. Time spent in the light compartment of the Dark-Light Box (left) (Two-way ANOVA, main effect of genotype F(1,29)=5.3, p=0.0287; n=17 wild-type, n=16 Mbd2−/−) and number of entries to the light side (right). Data are presented as mean ±SEM *p<0.05, **p<0.01 for wild-type vs *Mbd2*−/−. ### p<0.005 for wild-type over chance (50%) exploration (one-sample t-test). Note: no main effect for sex was observed in any of the tests. Therefore, behavioral data from males and females were collapsed within genotype for clarity, for full data see Fig S1–S3).

### Landscape of Mbd2 binding in the hippocampus

We focused our analysis on the hippocampus since the behaviors affected in our study are known to involve hippocampal functions [6–33]. In addition, other members of the MBD protein family were shown to have an important role in hippocampal function [13, 20, 34, 35]. We performed Mbd2 ChIP sequence on wild type and Mbd2−/− animals. We identified 2782 Mbd2 peaks annotated to 461 genes (with FDR<0.05). As expected, Mbd2 binds mostly to CpG-containing and GC-rich DNA regions (Χ^2^=123.49; df=1, p=2.2E-16 and Χ^2^=− 7168.2; df=1, p=2.2E-16, respectively). Many peaks were observed in proximity to transcription-start-sites (Fig2A). Interestingly, de-novo motif discovery found the transcription-factor E2F8 (p=1E-149) and NFAT5 (p=1E-108) to be highly enriched in Mbd2 binding peaks (Fig S5A). NFAT family members were also enriched in known motif-enrichment analysis (Fig S5B). The E2F family and NFAT family (activated by calcineurin) have been previously reported to have a role in neurogenesis, and brain development [36–39]. We also compared our data to publicly available ChIP-Seq data of the mouse hippocampus histone-marks [40]. As expected, Mbd2 bound overall to more repressive histone-marks (mostly H3K9me3) than to active histone-marks. However, a subset of the peaks was bound to both active and repressive histone-marks. This might imply that either Mbd2 binds sites that are bivalently marked in the same cell or that Mbd2 binds DNA regions that have different chromatin states in different cell populations within the hippocampus (Fig2B-C). Mbd2 co-occupancy with histone-marks on promoters, 3’ UTRs, TTS and exons was enriched for active histone-marks and depleted from repressive histone-marks consistent with an activating role (Fig2D). Exons and intergenic regions showed mildly enriched Mbd2 co-occupancy with repressive histone-marks (Fig2D). Pathway-analysis by GO-enrichment revealed enrichment of Mbd2 binding at several neuronal- and brain-related pathways such as: synapse assembly and axon guidance among others (Fig2E and FigS6). GO analysis of Mbd2 peaks associated with histone marks found that Mbd2 peaks associated with active and bivalent histone marks are enriched for brain-related pathways including: axon guidance, axonogenesis and modulation of chemical synaptic transmission thus further supporting a role for Mbd2 in neuronal development and function (Fig2F).

**Figure 2.**
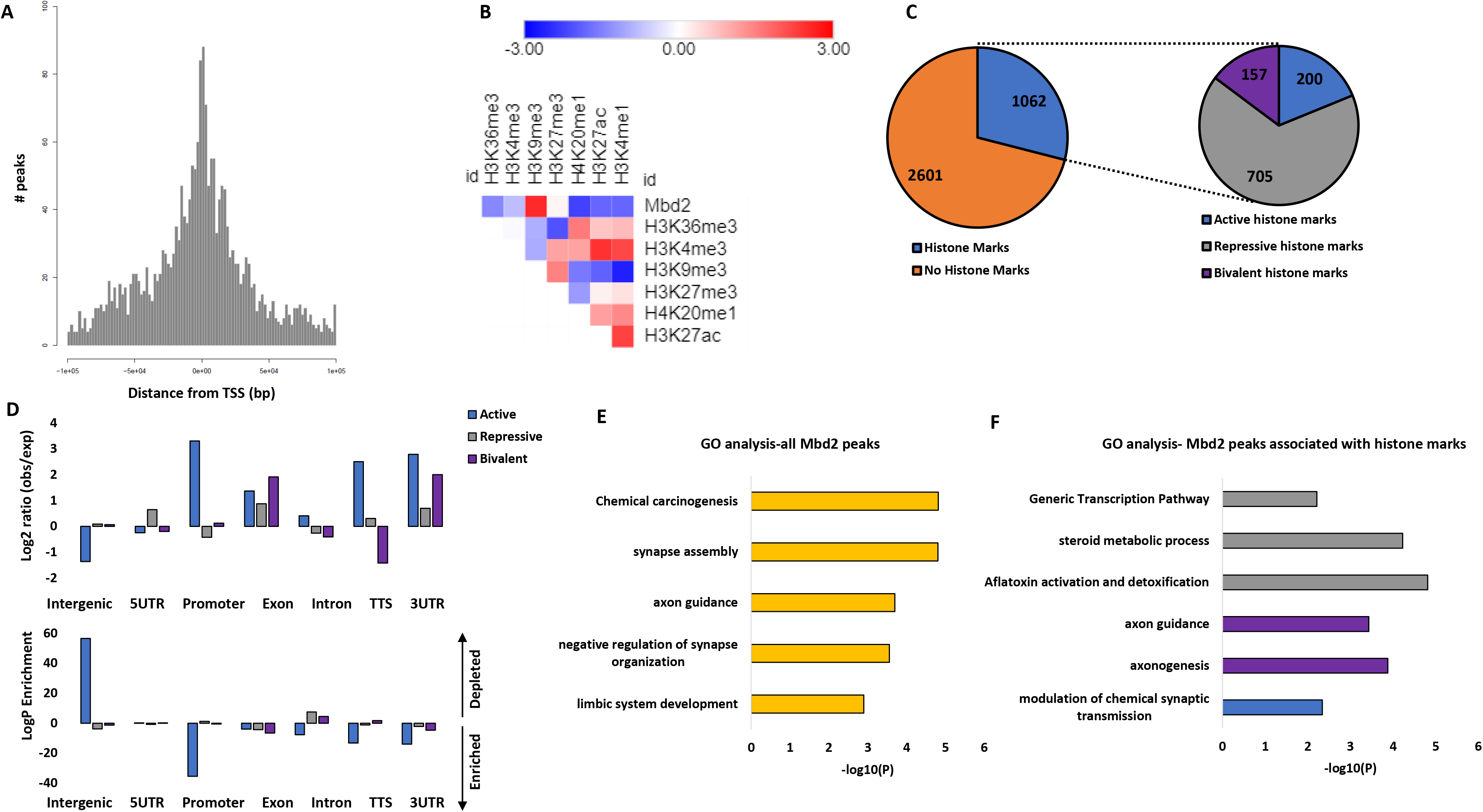
Landscape of Mbd2 binding delineated by ChIP-seq. A. Position of Mbd2 binding peaks relative to TSS. B. Heatmap of the overlap between Mbd2 peaks and the following histone-marks in the hippocampus: H3K4me1, H3K4me3, H3k2me3, H3K27ac, H39Kme3, H3K36me3 and H4K20me1. C. Analysis of the number of Mbd2 peaks located on histone-marks. D. Co-occupancy enrichment analysis (top) and significance level (bottom) of Mbd2 and histone-marks. E. Pathway analysis enrichment of Mbd2 binding peaks (Top 5 pathways are presented). F. Pathway analysis enrichment of Mbd2 binding peaks co-occupied with histone marks. TTS-Transcription Termination Site. UTR-untranslated region.

We selected several peaks of Mbd2 that were associated with promoters or enhancers in neuronal-related genes for further validation and for determining the impact of Mbd2 binding loss on transcription-initiation (see Table S1 for genomic coordinates of validated peaks). Quantitative-ChIP PCR confirmed binding of Mbd2 to these regions and loss of binding in *Mbd2*−/− mice (FigS7A). We then determined whether loss of Mbd2 binding in promoters or enhancers in *Mbd2*−/− mice affects transcription onset. *Mbd2*-deficiency resulted in reduced transcription onset since occupancy of RNA-polymerase phosphorylated on Serine5; (RNApolII(SP5)) the form found on promoters upon transcription-initiation [41, 42] was reduced in *Mbd2*−/− mice. Interestingly, this was associated with increased abundance of histone-mark (H3K4me1) marking enhancers in *Mbd2*−/− hippocampi (FigS7B-C). H3K4me1 modification marks active as well as inactive enhancers [43, 44]. H3K4me1 peaks that are flanking the TSS were shown to exhibit a peak-valley-peak pattern. The valley usually overlaps with transcription-initiation and RNApolII(SP5) peaks [45] reflecting a nucleosome-free zone and thus reduced histone presence and reduced signal for the histone-marks, which is prerequisite for transcription-initiation [46]. In transcriptionally inactive genes H3K4me1 peaks cover the entire TSS forming one continuous peak centered at the TSS and overlapping with the RNApolII(SP5) peaks. The increase of H3K4me1 concurrently with reduction of RNApolII(PS5) binding at the same position in *Mbd2*−/− mice is consistent with inhibition of transcription onset in response to *Mbd2* deficiency. These data suggest that Mbd2 is involved in activation of transcription turn on in these neuronal-specific promoters.

### Mbd2 is required for expression of neuronal genes

To further elucidate the role of Mbd2 in hippocampal gene-expression we delineated the transcriptomes of wild-type and *Mbd2*−/− mice using RNA-Seq. We found 2907 genes to be differentially expressed (FDR<0.05), of which 1590 genes were up-regulated, and 1317 gene were down-regulated in *Mbd2*−/− mice (Fig3A, and FigS8 for validations). Pathway-analysis by GO-enrichment found a robust down-regulation of neuronal-related pathways such as neuron-projection development, trans-synaptic signaling and behaviors (Fig3B and Fig S9A) which were highly organized in clusters based on clustering analysis of the GO-networks (Fig S9B). In striking contrast, the same analyses applied to up-regulated genes revealed mostly homeostasis and metabolism related pathways with no enrichment of neuronal-related pathways (Fig 3C and Fig S10A). GO-network analysis revealed poor clustering of the differentially expressed genes in these pathways (Fig S10B).

**Figure 3.**
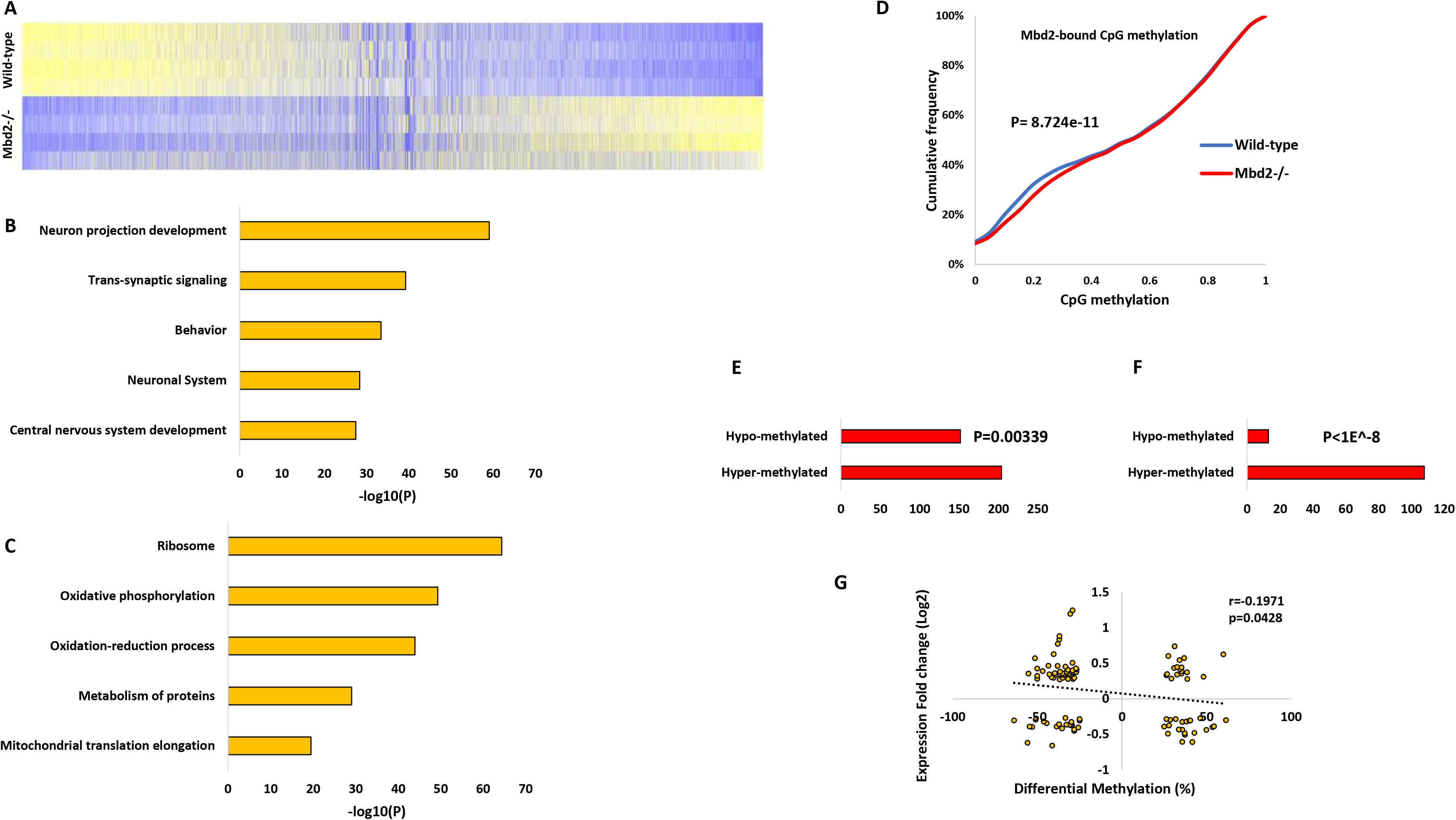
Effect of Mbd2 deficiency on gene expression and DNA methylation. A. Heatmap of all transcribed genes in *Mbd2*−/− and wild-type mice sorted by log2 fold-change. B. Pathway analysis enrichment of down-regulated genes (Top 5 pathways are presented). C. Pathway analysis enrichment of up-regulated genes (Top 5 pathways are presented). D. A cumulative distribution of methylation levels of regulatory DNA regions bound by Mbd2 demonstrating hyper-methylation in *Mbd2*−/− hippocampus (K-S test). E. Number of differentially methylated CpGs (with at least 10% change in methylation and p<0.001) in Mbd2-bound annotated genes showing that more CpGs became hyper-methylated then hypo-methylated (p=0.00339, binomial test). F. Number of differentially methylated CpGs (with at least 10% change in methylation and p<0.001) located within Mbd2-peaks (p<1E-8, binomial test). G. Pearson correlation between differential promoter DNA-methylation and gene-expression in *Mbd2*−/− hippocampi.

### The impact of *Mbd2* depletion on the DNA methylation landscape

MBD proteins are “readers” of DNA methylation and bind specifically to methylated DNA [4, 47]. However, it is possible that they also play a role in maintaining DNA methylation/demethylation landscapes as has been previously proposed. Previous studies in T-regulatory cells using *Mbd2*−/− mice have suggested that depletion of *Mbd2* causes hypermethylation of regulatory regions which in turn results in downregulation of gene-expression [48]. Therefore, we tested whether *Mbd2*-deficiency would alter the landscape of DNA methylation in the hippocampus. We mapped with targeted-bisulfite-sequencing the methylation state of regulatory DNA regions (promoters and enhancers, see methods) in the hippocampi of *Mbd2*−/− and wild-type mice at a single-base resolution. A global methylation analysis revealed that Mbd2 binding regions defined in wild-type mice by ChIP-seq were hypermethylated in *Mbd2*−/− mice (Fig 3D). Hypermethylation in response to *Mbd2* depletion was observed also in a genome-wide scale analysis which examined DNA outside Mbd2-bound regions (p<2.2E-16, K-S test, Fig S12C). The fact that changes in DNA methylation occur in regions that do not bind Mbd2 suggests *Mbd2*-deficiency could affect DNA methylation indirectly. We therefore examined our RNA-seq data for expression levels of methylated-CpG readers and modifiers. We found a significantly reduced expression of *Tet2, Dnmt3a, Mbd1* and *Mecp2* in *Mbd2*−/− mice (Fig S11). The reduced expression levels of enzymes which promote de-novo DNA methylation (*Dnmt3a*) and de-methylation (*Tet2*), together with the reduced levels of methyl-CpG binding proteins (*Mbd1* and *Mecp2*) likely contributed to changes observed in the landscape of DNA methylation in *Mbd2*−/− mice. Taken together these data suggest that loss of *Mbd2* affects DNA methylation in both directions. An analysis of the sequence properties of Mbd2-dependent DNA methylation suggests that *Mbd2*-deficiency results in increased methylation of low to intermediate methylated CpGs (10-40% methylation) but highly methylated CpGs (>40% methylation) are unaffected as expected if Mbd2 prevents hypermethylation. Partially methylated genes represent a group of genes that are heterogeneously methylated and possibly heterogeneously expressed across hippocampal nuclei. Mbd2 might be regulating the state of methylation of these genes.

We also analyzed differential-methylation at single CpG resolution. At this resolution differential-methylation analysis (at least 10% difference, FDR<0.05; Fig S12A) revealed 323 differentially-methylated CpGs, with 133 hypo-methylated CpGs (annotated to 103 genes) and 190 hyper-methylated CpGs (annotated to 119 genes) (differential-methylation data (p<0.001) is detailed at Table S2). This finding also supports a significant overall hypermethylation in *Mbd2*−/− mice hippocampus (p=0.0018, binomial-test). For pathway-analysis, we applied a more lenient significance cut-off for our data (p<0.001) resulting in 3005 differentially methylated CpGs (1519 hypo-methylated and 1486 hyper-methylated). Next, we focused the analysis on CpGs located on promoters (±1000bp from TSS). We found 494 hypo-methylated CpGs located in 460 gene-promoters and 476 hyper-methylated CpGs located in 434 gene-promoters (Fig S12A). Pathway-analysis for differentially methylated gene-promoters revealed adrenergic receptor signaling and sodium- and metal-ion transports as the top three hyper-methylated pathways (-log10p-values: 4.86, 4.16 and 3.42, respectively). In contrast, the top hypo-methylated pathways were not specifically related to neurons or brain functions, although neuronal ion-channel clustering and positive regulation of sodium ion-transport pathways were enriched (-log10p-values: 3.289 and 3.280, respectively) (Fig S12E-F). Overall, this analysis suggests an enrichment of hyper-methylated genes related to neuronal gene-pathways in *Mbd2*−/− mice hippocampi.

Next, we assessed how binding of Mbd2 affects single CpG DNA methylation in Mbd2-bound genes. We analyzed the changes in DNA methylation levels for CpGs (with at least 10% change and p<0.001) annotated to Mbd2-bound genes and found significantly more genes had hyper-methylated CpGs (P=0.00339, binomial test, Fig 3E). When analyzing only the subset of these CpGs which were located directly at Mbd2 binding peaks we found in almost all cases (~90%) hyper-methylation of these CpGs (p<1E-8, binomial test, Fig 3F). This is consistent with a role for Mbd2 binding in maintaining a hypomethylated state.

To assess the relation between promoter DNA methylation and gene-expression we determined the correlation between methylation and expression. We found a significant linear inverse correlation between promoter DNA methylation and gene-expression (Fig3G), supporting a role for DNA methylation in gene-repression in the hippocampus. Unexpectedly, a subset of differentially methylated CpGs in promoters showed positive correlation between expression and methylation. We explored this finding by analyzing DNA motifs around these CpGs. While CpGs with inverse correlation (n=66) had no significantly enriched motifs; CpGs with positive correlation (n=40) had 3 enriched motifs (p-values= 0.015-0.025). These enriched motifs were highly similar to known motifs of the KLF and SP transcription-factor families with top similarities to: KLF12, KLF14, SP3 and SP8 (Fig S12D). The KLF and SP transcription-factors are known to prefer binding to methylated-DNA (KLF) or to be unaffected by DNA methylation (SP) [49]. This finding suggests that genes whose methylation status does not correlate inversely with their expression, are regulated by transcription-factors that are not inhibited by DNA methylation.

We also validated our DNA methylation analysis results by targeted-sequencing of bisulfite-converted PCR amplicons of nine Mbd2 binding regions each of which annotated to a different gene (See Table S5 for delineation of the regions). Methylation levels in these regions strongly correlated with the levels obtained in the genome-wide capture sequencing analysis (range of Pearson’s r=0.884 - 0.913; Fig S13A-B). Loss of *Mbd2* promoted bi-directional alterations in DNA methylation status with enhanced hyper-methylation, as observed before (Fig S13C-D). Overall, there was a significant hyper-methylation in *Mbd2*−/− samples (Fig S13E) as was observed in the genome-wide capture-sequencing.

### Differentially expressed genes in *Mbd2*−/− mice are associated with differentially expressed genes in ASD individuals’ brains and ASD risk genes

Other MBD proteins were associated with ASD [12, 13, 34, 35, 50] and the presentation of behavioral abnormalities in *Mbd2*−/− mice described above resembles some of the observed symptoms of ASD. Therefore, we sought to explore the potential relation between our gene-expression findings and those found in post-mortem brains from neurodevelopmental disorders patients. We compared our data to the recently released cortex transcriptomic data from the psychENCODE consortium [51] for ASD, schizophrenia and bipolar disorder.

We found a highly significant overlap between differentially down- and up-regulated genes in Mbd2−/− mice and individuals with ASD (Fig 4A). We also found a weak, though significant, overlap between down-regulated (but not up-regulated) genes in Mbd2−/− mice and down-regulated gene in schizophrenia patients (Fig 4A). We did not find any significant overlap with bipolar disorder differentially expressed genes (Fig 4A). Using correlation analysis, we found a significant positive correlation of gene-expression fold-change levels between Mbd2−/− mice and ASD individuals but not between Mbd2−/− mice and schizophrenia or bipolar disorder patients (Fig 4B).

**Figure 4.**
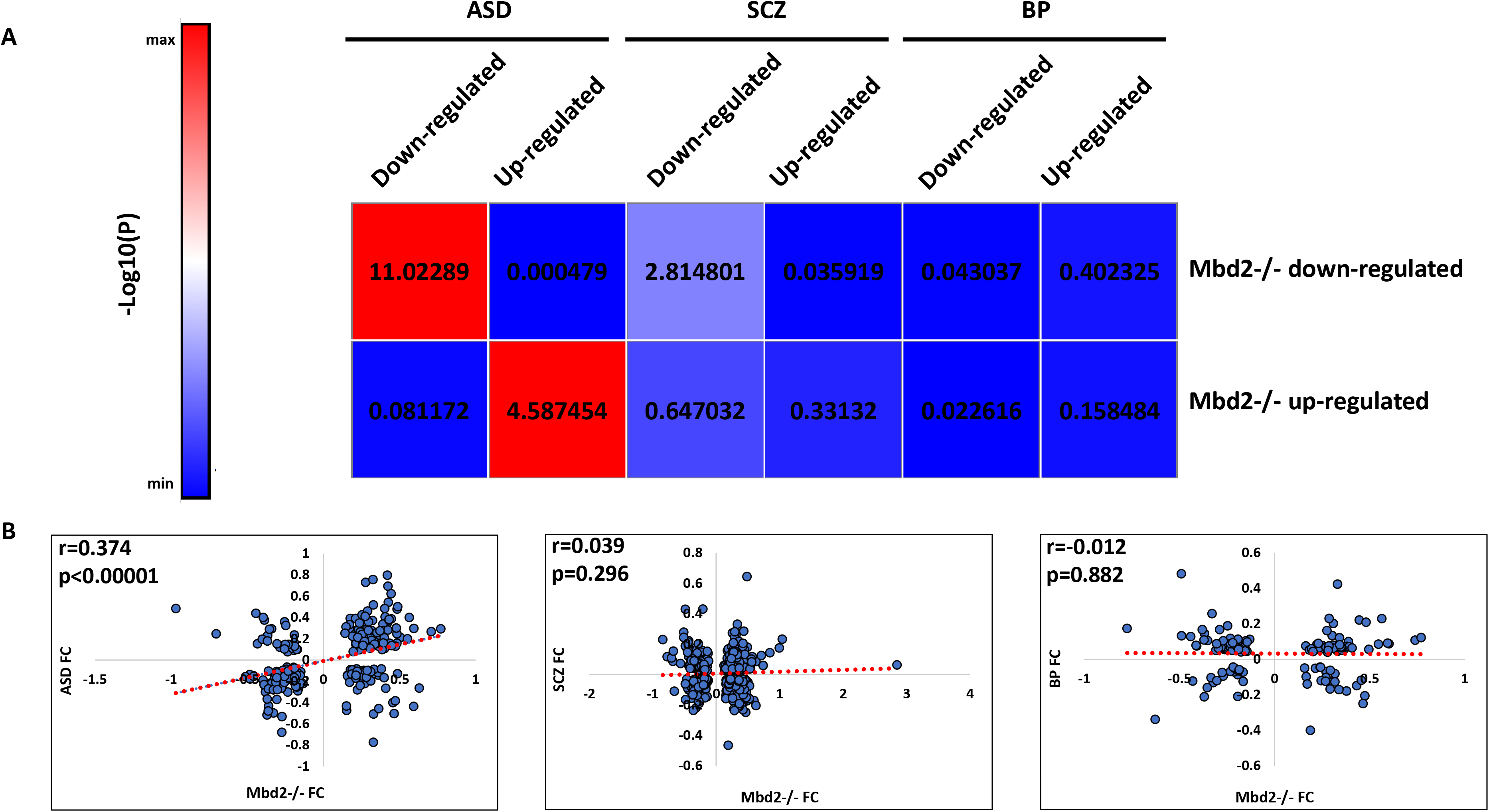
Overlap between *Mbd2*−/− mice hippocampal transcriptome and human brain transcriptomes in psychiatric disorders. A. Heatmap of log10(p-values) of differentially expressed gene overlap between *Mbd2*−/− mice and ASD, schizophrenia and bipolar disorder human brain. B. Pearson’s correlations of fold-changes in gene-expression levels between *Mbd2*−/− mice and ASD (n=286 genes), schizophrenia (n=686 genes) and bipolar disorder human brain (n=137 genes).

Next, we compared our gene-expression data with that of two human ASD risk-genes databases: a study by Sanders et al. [52]and the SFARI-gene database (see methods). We found a robust overlap between ASD-risk genes and down-regulated genes in *Mbd2*−/− mice. Twenty-four (~37%) out of 65 ASD risk genes (FDR<0.1) found by Sanders et al. [52] were down-regulated in *Mbd2*−/− mice (p=2.47E-12, hypergeometric-test, Fig5A and table S3). In contrast, only 3 genes from the 65 gene-list were up-regulated in *Mbd2*−/− mice (p=0.91 hypergeometric-test). The 24 down-regulated genes common to mice and humans also formed a significant protein-protein interaction network (p= 8.12E-6, Fig S14A) with some of the top GO pathway annotations related to “regulation of biological and cellular processes” and “nervous system development” (Table S4).

Next, we compared our RNA-seq results to the SFARI-gene list of ASD-associated genes which contains all known human genes associated with ASD. Here again, out of 951 human genes associated with ASD, 125 (13.14%) were down-regulated (p=8.8E-13, hypergeometric-test, Fig 5B and table S3) while only 44 were up-regulated in *Mbd2*−/− mice (4.62%, p=0.99, hypergeometric-test, Fig 5B and table S3). Analysis of Mbd2-regulated ASD-associated gene-lists revealed significant protein-protein interactions enrichments. The 125 down-regulated genes which appears also in the SFARI ASD-associated gene-database projected into a significantly enriched protein-protein network (p<1E-16; Fig S15A) with “nervous-system development” and “modulation of excitatory postsynaptic potential” as the most enriched GO-pathways within the network (p=5.84E-15 and p=2.14E-13 respectively, table S4). The 44 up-regulated genes which appear also in the SFARI ASD-associated gene-database projected into a significantly enriched protein-protein network (p=0.00559; Fig S15B) with “dendritic spine morphogenesis” and “modulation of excitatory postsynaptic potential” as the most enriched GO-pathways within the network (p= 0.00162 in both cases, table S4).Taken together, these transcriptomic and genomic cross-species comparative analyses suggest that Mbd2 might serve as an up-stream regulator to many ASD-associated genes. This is in line with previous observations that showed regulatory roles for Mbd2 in liver, breast and prostate cancer genes in cancer cell lines [6–55], NGFIA in hippocampal cells [19] and *Foxp3* gene [48] in regulatory T-cells. We further analyzed the protein levels of few members of the protein-protein network of ASD-risk genes and found reduced protein levels of these genes in *Mbd2*−/− mice (Fig 5C) which agrees with the mRNA reduction we observed before.

**Figure 5.**
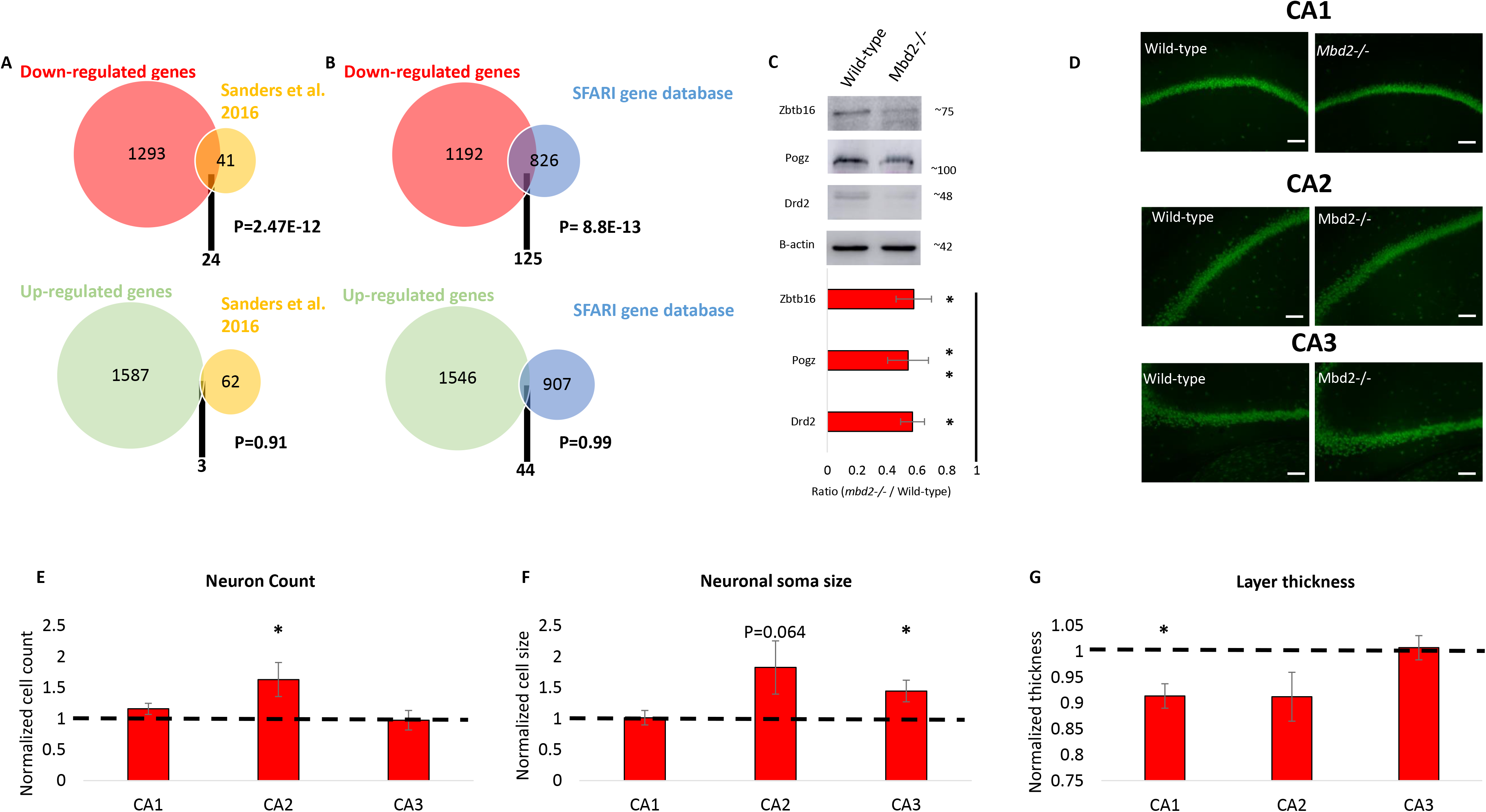
Overlap between down-regulated genes in *Mbd2*−/− mice and ASD-risk genes. A. An overlap analysis revealed a significant intersection between human ASD high risk genes and down-regulated (Top) but not up-regulated (Bottom) genes in *Mbd2*−/− hippocampi. B. An overlap analysis revealed a significant intersection between human ASD risk genes based on the SFARI Gene Project and down-regulated (Top) but not up-regulated (Bottom) genes in *Mbd2*−/−. (Hypergeomtric tests). C. Western blot analysis showing reduced protein levels of Drd2, Pogz and Zbtb16 in *Mbd2*−/− mice hippocampi. Proteins weights as determined using molecular weight markers are indicated. D. Representative immunofluorescence images of the hippocampus (CA1-CA3 layers) in wild-type and *Mbd2*−/− mice showing neuronal nuclei immunoreactivity (NeuN, green immunoreactivity). Scale bar: 100μm. E. Quantification of neuronal cell count in hippocampal layers. F. Quantification of average neuronal soma size in hippocampal layers. G. Average thickness of hippocampal layers.

Morphological changes in the hippocampus were observed as well. We found increased neuron count in the CA2 region of the hippocampus (Fig 5E and Fig S16), increased neuronal soma size in CA3 (Fig 5F) and reduced CA1 thickness (Fig 5G). We also found a negative correlation between neuron soma size and neuron count (for CA1 and when all regions data was collapsed) and between neuron soma size and thickness (for CA2). These changes indicate an increased neuronal density and reduced thickness in the hippocampus of *Mbd2*−/− mice, phenomena which were previously reported in ASD rodent models and human subjects [6–58]. Increased neuronal density was also observed in the cortex of children with autism [59].

### Hippocampal Mbd2 down-regulation impaired cognitive and emotional behaviors

Next, we asked whether hippocampal Mbd2 is casually involved in regulation of behaviors which are impaired in adult *Mbd2*−/− mice. We therefore examined whether Mbd2 down-regulation in adult mice affects some or all of the behavioral changes seen in *Mbd2*−/− mice which lacked the gene both during embryogenesis and post-natal development. To this end, we constructed several shRNA lentiviruses which targeted *Mbd2* mRNA (sh-*Mbd2*; Fig S17A-B). The construct which was the most active in *Mbd2* depletion as tested *in-vitro* was infused *in-vivo* into the hippocampus of adult wild-type mice to knockdown Mbd2 in-vivo (Fig 6A-B). GFP-expressing-cells were found near the injection site and up to the dentate gyrus and stratum radiatum of CA3 (Fig S17C). Following the infusion and recovery we tested the mice in the same behavioral tests that were found to be abnormal in *Mbd2*−/− mice. Down-regulation of Mbd2 did not affect locomotion in an open-field (Fig 6C) but increased self-grooming time and average grooming bout duration (Fig 6D-F). Interaction time in the social-interaction test was not affected by Mbd2 down-regulation (Fig 6H). However, Mbd2 down-regulation resulted in failure of treated mice to remember location of objects in the object-location-memory test. While control mice explore the object in the novel-location significantly more than the familiar location (p<0.05, one-sample t-test), sh-*Mbd2*-treated mice did not explore the object in the novel location any longer than chance level (p>0.05, one-sample t-test, Fig 6G). In the Dark-Light-Box test, sh-*Mbd2*-treated mice showed increased anxiety expressed as reduced time in the light compartment (Fig 6I-J). Taken together, these results show that knockdown of hippocampal Mbd2 in adult mice recapitulate most of the behavioral abnormalities found in *Mbd2*−/− mice but not the social interaction deficiency.

**Figure 6.**
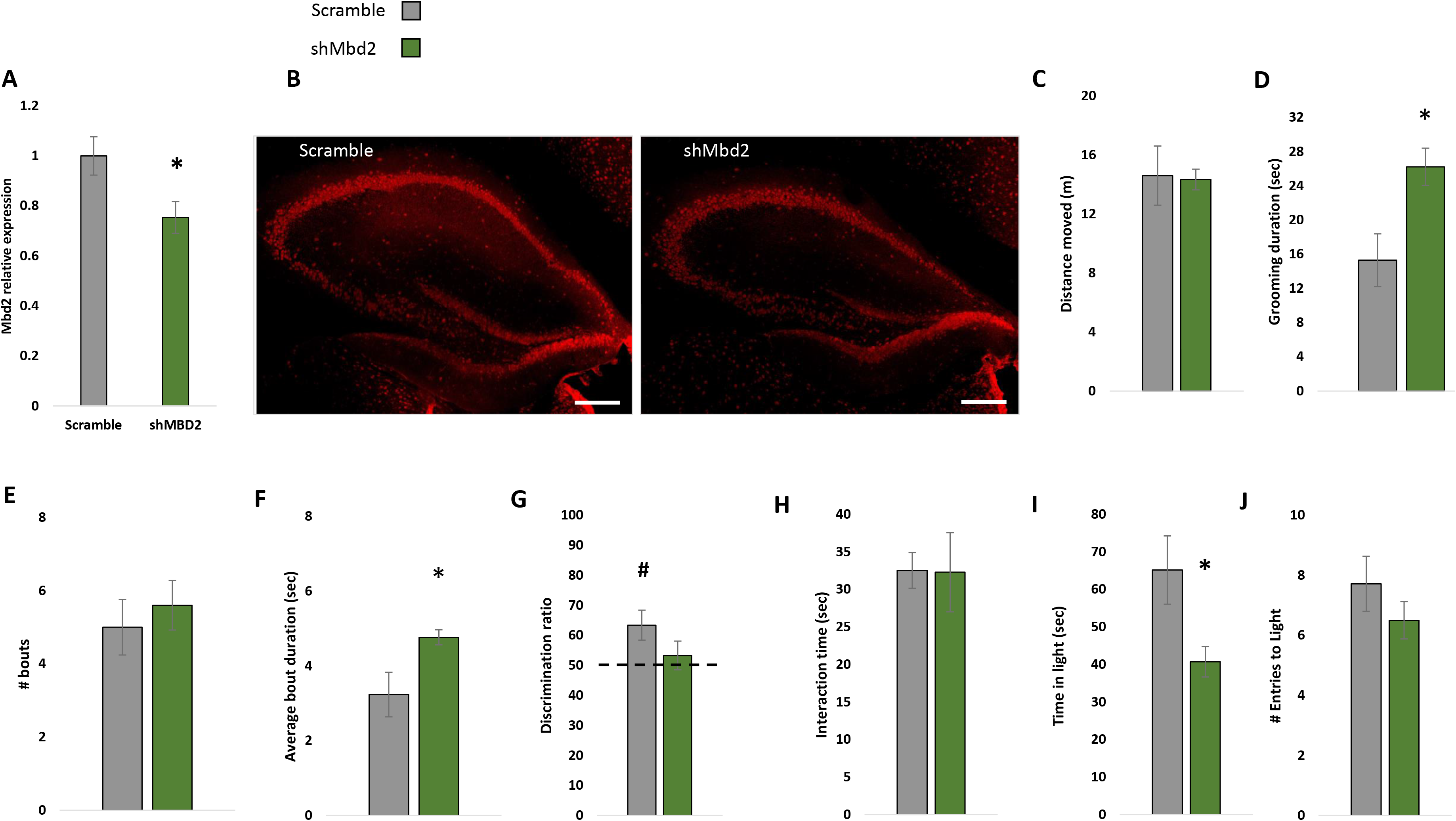
Virus-mediated Mbd2 down-regulation. A. shMbd2 viral construct reduces *Mbd2* mRNA. Mbd2 mRNA levels were reduced in the hippocampus from shMbd2-infused mice compared with scramble control-infused mice. Scramble n=6, sh*Mbd2* n=4 (T test). B. Representative micrograph of Mbd2 immunolabeling in hippocampal slices from Scramble- and sh*Mbd2*-infused mice. Scale bar: 200μm. C. Locomotion in Open-field box. D. Self-grooming during open field. E. Number of self-grooming bouts. F. Average self-grooming bout duration. Discrimination ratio in object-location memory test. H. Social interaction test. I. Time in the light compartment of the Dark-Light Box (left) and number of entries to the light side (right). Data are presented as mean ±SEM *p<0.05, for wild-type vs Mbd2−/− (t-test). # p<0.05 for wild-type over chance (50%) exploration (one-sample t-test). Scramble n=7, shMbd2 n=6.

## Discussion

There is growing evidence for the crucial role of epigenetic mechanisms in neuropsychological disorders and CNS function. In this study, we tested the role of Mbd2, a methylated-CpG binding protein, in gene-expression and brain function. Mbd2, like other MBD proteins, serves as a “reader” of the epigenome [4] translating DNA methylation marks into gene-expression regulation mechanisms. It has previously noted that loss of Mbd2 results in behavioral deficits such as impaired maternal care, poor nest building and mild memory impairment [24, 25]. Here we report that loss of Mbd2 results in cognitive, social and emotional deficits. Taken together these results imply a role for Mbd2 in normal brain function. Most of the phenotypic findings were replicated by hippocampus-specific down-regulation of Mbd2 in wild-type mice (increased self-grooming, impaired memory-retention in the object-location memory test, and anxiety-like behavior in the dark-light box) providing further evidence for the causal role of hippocampal Mbd2 in regulating behavioral phenotypes including long-term spatial memory and emotional control. Social interaction was not affected by *Mbd2* knockdown. It is possible that the level of inhibition of Mbd2 achieved by lentiviral knockdown was insufficient to affect social interaction. Alternatively, hippocampal Mbd2 might play a developmental role in social interaction behaviors and might not be required for maintenance of this behavior in adults.

Since the effects of *Mbd2*-deficiency on behavior were broad and related to hippocampal function [6–33], we used genome-wide approaches to define the landscape of Mbd2 binding in the hippocampus and then determined how its deficiency affects the landscapes of DNA-methylation and gene-expression using *Mbd2*−/− mice. Consistent with an important role in brain function as detected by the behavioral assays presented here, pathway-analyses revealed a highly clustered and networked enrichment of genes relating to cognitive functions and brain development such as trans-synaptic signaling, synapse organization and behavior.

Mbd2 regulates gene-expression by binding to methylated CpGs in DNA [60]. MBD proteins are generally considered to have a repressive role on gene-expression by interacting with chromatin modification inactivating complexes [61]. However, evidence suggests other MBD proteins are involved in gene-expression in more than one way. For example, hypothalamic Mecp2 dysfunction led to robust changes in gene-expression with 85% of the genes activated by Mecp2; a process possibly mediated by the interaction of the transcriptional-activator Creb1 with Mecp2 in binding gene-promoters and regulatory regions [62]. Similarly, Mecp2 promotes the expression of *Fopx3* in regulatory T-cells (Treg), a key regulator of Treg function, by collaborating with Creb1 [63]. Neuronal Mbd1 was also found to have a mixed effect on gene-expression in the hippocampus with more than half of the genes down-regulated in *mbd1−/−* mice, many of them associated with neurogenesis [23]. Mbd2 was previously shown to be involved in both suppression of promoters through recruitment of transcriptional-repressors, and in activation of promoters in cancer cells and the hippocampus through recruitment of transcriptional-activators [19, 53–55] such as CBP and NGFIA [19, 55, 64] as well as by targeting DNA demethylation [65, 66] possibly through recruitment of oxygenases such as Tet2 [48].

Consistent with these data we found evidence for the bimodal function of Mbd2 in the hippocampus. *Mbd2*-deficiency resulted in both gene activation and repression as well as hypermethylation and hypomethylation. Interestingly, the neuronal specific functions seem to be mainly repressed and hypermethylated by *Mbd2*-deficiency, suggesting a mostly activating role for Mbd2 at neuronal-specific genes. This is supported by the observation that Mbd2 localizes to promoter and transcription start sites.

To further examine the role of Mbd2 in transcriptional activation/repression, we examined several promoters and enhancers that bind Mbd2 and examined the consequences of *Mbd2*-deficiency on transcription-initiation. Most of the genes examined had reduced RNApolII(PS5) occupancy and increased H3K4me1 masking of the transcription-initiation region indicating reduced transcription-initiation upon *Mbd2*-deficiency.

Both genetic and environmental factors are believed to be involved in the etiology of ASD and other neurodevelopmental disorders. DNA methylation has been proposed as an epigenetic interface which links environmental factors to genomic susceptibility in ASD [6–73] and neurodevelopment [74, 75]. MBD proteins as readers of DNA methylation are therefore potential mediators which translate the altered DNA methylation landscape into gene-expression and ultimately behavioral changes. While the involvement of MBD2 in psychiatric disorders was previously suggested [14, 16, 76], our study is the first to provide a possible mechanism. Interestingly, the list of genes that we found here to be Mbd2-dependent was significantly enriched in ASD-associated genes according to brain transcriptomic and GWAS studies [51, 52]. Moreover, the behavioral phenotypes that we and others have characterized in Mbd2−/− mice resemble several of the phenotypes frequently observed in ASD individuals such as social-avoidance, repetitive behavior, anxiety and deficits in learning and memory. This is accompanied by morphological changes in the hippocampus similar to those previously reported in ASD individuals and ASD rodent models such as increased number of neurons [56, 59] accompanied by reduced thickness of hippocampal sub-regions [57, 58]. It is therefore possible that *Mbd2*−/− mice have impaired neuronal development which results in an increased number of presumably immature neuron but less mature neurons. This pattern has been described before in *Mbd1−/−* mice [13, 23].

Taken together, our findings further highlight the importance of MBD proteins in neuropsychological disorders. Our findings also provide evidence that Mbd2 is a regulator of neuronal gene-expression, brain morphology and behavior.

## Methods

### Animals

The *Mbd2*−/− line was created in the lab of Dr. Adrian Bird [24]. The embryos of heterozygous *Mbd2(−/+)* mice were kindly gifted by Dr. Brian Hendrich from the Department of Biochemistry, University of Cambridge which were bred at McGill University animal facility. Mice were housed in groups (3-5/cage) with 12-h light/dark cycles under conditions of constant temperature (23°C) and humidity (50%), and free access to food and water. All procedures were carried out in accordance with the guidelines set down by the Canadian Council of Animal Care and were approved by the Animal Care Committee of McGill University.

Previous studies [24] found that *Mbd2*−/− dams show impaired maternal care and nurturing behavior, which in turn results in slower weight gain of the pups. These phenomena were not observed in heterozygote females and fathers genotype had no effect on weight gain. Cross-fostering experiments demonstrate pups’ normal weight can be rescued by fostering with wild-type dams and impaired by fostering with Mbd2−/− dams [24].

Therefore, all littermates were bred from heterozygote parents to avoid confounding effect of maternal behavior. All ex-vivo experiments were conducted on 10-12 weeks old mice.

### Behavioral procedures

All behavioral experiments were conducted on 10-12 weeks old mice.

#### Open-Field and Self-Grooming

Mice were placed in an open field (45×45 cm) apparatus with 30-cm-high walls. Their locomotor activity, exploration and self-grooming behaviors were measured for 5 min. The open-field test was also used as a habituation session for the object-recognition and object-location tests.

#### Object-Location Memory (OLM)

Each mouse was placed for 5 min in the open field with two identical objects (A1 and A2) positioned in two adjacent corners. Exploration was defined as sniffing or touching the object with the nose, whiskers and/or forepaws. The exploratory preference for each object was calculated as the percent time (t) spent exploring that object relative to the total time spent exploring both objects [(tA2/(tA1+tA2) x100]. For the long-term memory (LTM) test conducted 24h after the training session, the same mouse was allowed to explore the field for 5 min in the presence of the same objects in the same settings with the exception that one of the objects was displaced to a novel location. Visual cues (adhesive tape) were placed on two adjacent walls as described before [31].

#### Social Interaction Test

The test was performed as described before [28]. Previous study did not find changes in social behavior in the three-chamber social approach test in Mbd2−/− mice [25]. Therefore, we selected the social interaction test based on reports suggesting its increased sensitivity over the three-chamber social approach test [6–28]. The duration of the test was 5 minutes. Sniffing, close following, and allo-grooming were defined as social interaction. No aggressive behaviors were found during the tests.

#### Spontaneous Alternations in Y Maze

The Y maze was a standard 30 × 6 × 15 cm (Length X Width X Height) grey apparatus. Mice were allowed to freely explore the maze for 5 minutes. An entry to an arm was considered when all four paws were inside the arm. The measures included spontaneous alteration performance (i.e., a successful triad), alternate arm returns (AAR) and same arm returns (SAR). Total number of entries was assessed as well.

#### Dark-Light-Box

Anxiety-related behavior of the mice was tested in the Dark-Light Box consisting of two chambers connected by an open gate. Mice were placed near the gate in the light chamber facing the dark chamber. Latency to first entry to the light side, number of exits and total time in the aversive light compartment were recorded for 5 minutes.

We analyzed all the behavioral data for male and female mice and did not find any significant differences in behavior across sexes (two-way ANOVAs, p>0.05 for main effect of sex and for interactions in all cases, see also FigS1 and FigS2). We therefore grouped male and female hippocampi for all the following experiments with balanced numbers of male and females between groups.

### Lentivirus Production

Three *Mbd2* shRNA plasmid were purchased from Dharmacon™ (Lafayette, CO). The day before transfection, 1× 106 HEK293T cells were seeded in 10cm plate (DMEM medium supplemented with FBS and L-Glutamine, (Gibco, Ottawa, ON)). Next day, a 3 rd generation lentivirus was assembled by co-transfection of 4 vectors (5 μg each): pGIPZ vectors with one of three shRNA inserts or scramble sequence, a VSV-G envelope-expressing plasmid pMD2.G (Addgene plasmid #12259), and the packaging plasmids: pRSV-Rev (Addgene plasmid #12253) and pMDLg/pRRE (Addgene #12251). shRNAs sequences were: sh-*Mbd2*-1: 5’-AATTTGTTCTGTTACATCT-3’; sh-*Mbd2*-2: 5’- TTTCGGATCACTTCCTCCT-3’; sh-*Mbd2*-3: 5‘-TCTTCTGTAATTTACTAGG-3’; scrambled: 5’- GCCUUGGCAGCCUAGGCGA-3’. Lentivirus assembly was performed in the following manner: all four plasmids were mixed in 50 μl of OPTI-MEM (Invitrogen, Ottawa, ON). 45 μl of Fugene HD transfection reagent (Roche, Laval, QC) was diluted in 400 μl of OPTI-MEM. Plasmid mix was added to the diluted Fugene HD, mixed gently by pipetting and incubated for 30 min. at room temperature. Meanwhile, HEK293T cells were washed with PBSx1 and replaced with appropriate medium. The Fugene HD-DNA mix was added to the cell medium. After 48 hours of transfection of four plasmids, medium, containing virus was collected and passed through 0.45um filter.

### *In-Vitro* Mbd2 Viral-mediated down-regulation

In order to examine the ability of sh-*Mbd2* to down-regulate Mbd2 expression NIH3T3 cells were infected with the three different sh-*Mbd2* lentiviruses. At 72 h after infection, cells were lysed, and the RNA extracted using Trizol according to the manufacturer’s protocol. RNA was reverse-transcribed to cDNA using M-MuLV Reverse Transcriptase kit (NEB, Ipswich, MA) and cDNA was analyzed with QPCR.

### Stereotactic Surgery and Viral Infusion

Eight-week-old mice were anesthetized with a Ketamine/Xylazine mixture and placed in a stereotaxic apparatus (Kopf Instruments, Tujunga, CA). Bilateral infusions of a shRNA-*Mbd2*-containing lentivirus (sh-*Mbd2*-1, or scramble control (n=9/group) were performed with a borosilicate glass capillary lowered to the hippocampus. Coordinates relative to bregma were: AP −1.9 mm, ML ±1.25 mm, and DV −1.75 to −2.0 mm; based on previous reports [77]). Infusion was done with Auto-Nanoliter Injector (Drummond Nanoject II, Broomall, PA) at a rate of one 38 nanoliter infusion (lasts for 2 sec) followed by 20 seconds interval for a total of 2μl/hemisphere. Glass capillary was remained in place for at least 2 minutes after the infusion to avoid reflux. Carprofen was injected post-surgery for at least 72h. Animals had 2 weeks recovery period before behavioral testing. Animals which had poor recovery or high levels of distress were euthanized according to the guidelines set by the Animal Care Committee of McGill University. We therefore performed the behavioral tests on 7 scramble-infused and 6 shMBD2-infused mice.

### Chromatin-Immunoprecipitation Followed by Sequencing (ChIP-Seq)

For all ex-vivo experiments, mice (10-12 weeks of age) were sacrificed, the brains were rapidly removed, and bi-lateral hippocampi were isolated, flesh frozen and stored at −80°C for later analysis. For Chromatin-Immunoprecipitation, two independent cohorts of wild-type and two independent cohorts of Mbd2−/− mice hippocampi (n=6-8) were pooled, homogenized in 1 X PBS including 1% formaldehyde, and the homogenates were kept for 10 min at 25°C. Cross-linking reactions were stopped by the addition of glycine (125 mM) for 10 min at 25°C. Fixed chromatin samples were then homogenized in cell lysis solution (PIPES 5 mM (pH 8), KCl 85 mM, NP40 0.5%) and centrifuged for 5 min at 3000 rpm, 4°C. Pellets were resuspended in RIPA-light solution (NaCl 150 mM, SDS 0.3%, Tris-HCl 50 mM (pH 8)) and sonicated using a Covaris E220. Sonicated chromatin samples were then centrifuged for 15 min at 14000 rpm, 4°C. Pellets were resuspended in 1 ml of RIPA-light solution. Chromatin samples were pre-cleared with 50 μl of Dynabeads protein G (Life Technologies, Ottawa, ON) pre-blocked with BSA and incubated overnight at 4°C with an anti-Mbd2 (IP grade, Epigentek-A1007, Burlington, ON) antibody. Input control was treated identically the same way except for not adding an anti-Mbd2 antibody to the solution. Antibodies and chromatin were then mixed with 100 μl of Dynabeads protein G for 3 hours at 4°C. The beads were then washed with RIPA-light solution, then with wash-solution (Tris-HCl 100 mM (pH 8), LiCl 500 mM). Protein–DNA complexes were eluted from the beads, de-cross-linked, treated with proteinase K and purified. The DNA concentration was determined by fluorometry on the Qubit system (Invitrogen, Ottawa, ON). A total of 10–12 ng DNA were used for the preparation of the libraries. The immunoprecipitated DNA and input DNA were sheared a second time with the Covaris E220 instrument in 53 μl reaction volume (duty factor 10%, Pic Incident Power 175, Cycles per burst 200, time 360 sec) to obtain fragments in the size range of 150 bp followed by purification with AMPure XP beads (×1.8v/v) (Beckman Coulter A63881, Indianapolis, IN). Purified DNA was resuspended in 45 μl elution buffer. Libraries of the chromatin immunoprecipitated and input DNA fragments were prepared using the Tru Seq DNA Low Throughput Protocol (Illumina). PCR enrichment of ligation products was performed using the Illumina Primer Cocktail; 15 cycles of PCR were performed for ChIP libraries and 10 cycles for the input. The libraries were purified using AMPure XP beads ×1.0 v/v. Quality of libraries was validated by 260 nm absorbance measurement, quality control on HSdna chip (Agilent Bioanalyzer: size of libraries around 275bp) and quantification by Q-PCR with Kappa Library Quantification kit for Illumina Sequencing Platforms (KAPPA Biosystems). The DNA concentration of the different sequencing libraries was from 40 to 500 nM. Clusters (13.5 pM) were generated using TruSeq PE Cluster Kit v3, for cBot protocol, which was followed by 50bp either single or pair--end sequencing, on an Illumina HiSeq 2000, per the manufacturer recommendations.

### Bioinformatic Analysis of the ChIP-Seq Data

Read quality for all next-generation sequencing experiments was assessed using Fastqc (http://www.bioinformatics.babraham.ac.uk/projects/fastqc/) which confirmed high read quality and inconsequential levels of adapter contamination. Reads were aligned to the mouse reference genome (mm9 assembly) using Bowtie [78] with default parameters. Low quality alignments and alignments for read pairs with multiple possible genomic alignments were omitted. Duplicate read pairs were removed. Read count peaks corresponding to likely binding sites were identified for each sample by MACS2 [79] with callpeak -nolambda -q 0.05 parameters as described before [80], input DNA reads were subtracted from the Mbd2 binding to control for nonspecific peaks. Peaks with false discovery rates less than 0.05 were selected as high-confidence binding sites. Sites were annotated with their genomic locations relative to nearby genes using HOMER with default parameters. De-novo motif discovery was done with HOMER with background adjusted to CpG context and the default cumulative binomial distribution test.

### Q-ChIP

Hippocampi from 5-6 mice per group as biological replicates for validation of ChIP-seq identified Mbd2 peaks were subjected to the same ChIP protocol used for the ChIP-Seq. To exclude the possibility of idiosyncrasy of the anti-Mbd2 antibody used in the ChIP-seq experiment, we used another antibody for this experiment (Imegenex, IMG147, Oakville ON) and IgG antibody (Santa Cruz, Mississauga, ON) served as an additional control. For RNApolII(SP5) and H3K4me1 ChIP experiments antibodies from Abcam were used (ab5408 and ab8895, respectively. Toronto, ON). Purified DNA was resuspended in 40ul elution buffer. For QPCR analysis, SYBR green quantitative PCR was performed using the LightCycler^®^ 480 system (Software 3.5, Roche Molecular Biochemicals). To determine the relative enrichment, the 2 - ΔΔCt method was used with normalization to IgG and input data.

### RNA-Sequencing and Data Analysis

DNA and RNA from 8-10 mice was isolated and purified with AllPrep-DNA/RNA/miRNA-universal kit (Qiagen, Montreal, QC) and concentrations were determined by fluorometry on the Qubit system (Invitrogen, Ottawa, ON). Ribosomal RNA was removed using the Ribominus kit (Invitrogen). cDNA libraries (4 per group) were generated and sequenced using an Illumina Hiseq 2500 (100 bp pair-end runs), as instructed by Illumina’s RNA-seq protocols. Reads were deduplicated as described above. Reads were aligned to the mouse reference transcriptome (mm9) using the STAR aligner [81]. Differential expression was analyzed using DeSeq2 with an FDR of 0.05 [82].

### Human Brain Transcriptomic Data Analysis

The PsychENCODE Consortium (Gandal et al.2018) recently published RNA sequencing data from brain samples of 1695 individuals with ASD, schizophrenia, bipolar disorder and controls. We used this published database to create gene-lists of known human genes which were differentially expressed (up- or down-regulated) in human brains (FDR threshold<0.05). Overlaps between these human genes and differentially expressed genes in Mbd2−/− mice hippocampi were analyzed per disease and by direction of difference from healthy controls using hypergeometric test. Next, we analyzed the correlations between the magnitude of changes (expressed as Fold change) in gene-expression between-species for each disorder.

### Human Genetics Data Analysis

ASD risk genes list was extracted from Sanders et al. [52] which integrates genomic data from several large recent studies (total n = 10,220 individuals from 2,591 families). The data include 65 risk genes (FDR<0.1). In addition, data from the SFARI GENE project (gene.sfari.org) were extracted. This database contains gene-list of ASD associated genes based on human and animal model studies. Genes are grouped according to criteria for the strength of evidence for each gene (a total of 970 genes as of 23 November 2017). We excluded 19 genes categorized in category 6- for genes studied in human cohorts and findings suggest against a role in autism, having a total of 951 ASD-associated genes from this database. Next, we assessed the overlap between ASD risk genes from each of the databases to the ortholog mouse genes which were differentially expressed in *Mbd2*−/− mice. Significance was determined by a hypergeometric test.

### Quantitative polymerase chain reaction (qPCR)

RNA was extracted from 6-9 mice per group as biological replicates for validation of RNA-seq cDNA was prepared with random hexamer primers (Invitrogen, Ottawa, ON) and a reverse transcription kit (NEB, Ipswich, MA) according to the manufacturer protocol. *Actb* was used as the reference gene. SYBR green quantitative PCR (qRT-PCR) was performed using the LightCycler^®^ 480 system (Software 3.5, Roche Molecular Biochemicals). To determine the relative concentration of mRNA expression, the 2 −ΔΔCt method was used.

### Capture bisulfite sequencing and DNA methylation mapping

DNA was extracted from two cohorts of mice (8-10 mice/group per cohort). Each cohort was pooled and sequenced as detailed below. The SeqCap Epi Enrichment System (Roche NimbleGen, Laval, QC) for targeted bisulfite sequencing of promoters and enhancers in the mouse genome was used exactly as we described before [83, 84]. Mouse target probes (mm9) were custom designed based on H3K4me1 and H3K4me3 signals from mouse public ChIP-seq data. Biotinylated target probes were designed for both strands of bisulfite converted genomic DNA. Bisulfite treated genomic DNA was ligated to methylated next generation sequencing (NGS) adaptors, hybridized to the biotinylated oligonucleotide probes followed by a series of washes of off-target DNA sequences and unbound DNA. Isolated DNA then underwent PCR amplification and sequenced on Illumina Hiseq 2000 with pair-end 50bp reads and a technical repeat with 125bp pair-end reads was performed as well.

### Analysis of Differentially Methylated Cytosines

Sequences were aligned to the mm9 mouse reference genome using Bsmap v2.89 [85]. Output data were strand-sorted, filtered, and deduplicated with Picard tools. Next, methylation levels and coverage levels were extracted with methratio.py command in Bsmap. Differential methylation cytosines were analyzed with methylKit R package [86] with FDR threshold of 0.05 and at least 10% difference in methylation levels. Differentially methylated positions were annotated with HOMER [87]. De-novo motif discovery for differentially methylated promoters was done with RSAT [88]. We created 200bp sequences, each centered at the differentially methylated CpG as input for this analysis and performed RSAT motif discovery followed by comparison to known motifs from JASPER Core Nonredundant Vertebrates database.

### Targeted Sequencing of Bisulfite-Converted PCR Amplicons

To validate the results obtained from the genome-wide capture bisulfite sequencing we used an independent cohort of mice (5 wild-type, 3 *Mbd2*−/−). Hippocampal DNA was extracted as described above with AllPrep-DNA/RNA/miRNA-universal kit (Qiagen) and bisulfite-converted with EZ DNA Methylation-Gold Kit (Zymo Research). PrimerSuite software [89] (http://primer-suite.com/) was used to design PCR primers specific for bisulfite-converted DNA for genomic loci which we found to have Mbd2 binding peaks (see table S5 for chromosomal positions and gene names for the amplicons). PCR products were cleaned with a Qiagen kit, and libraries were prepared with NEBNext^®^ Ultra™ II (NEB). Libraries were sequenced on Illumina MiSeq with pair-end 250bp reads. Analysis of amplicons DNA methylation was done as described above for the genome-wide capture bisulfite sequencing.

### Protein-Protein Network Analysis

Differentially expressed genes which were identified also as ASD-associated genes, were analyzed for protein-protein association. We used STRING v10.5 [90] for this analysis with default settings (medium confidence threshold 0.4).

### Tissue Processing

Mice were anesthetized before trans-cardiac perfusion with ice-cold saline solution (pH 7.4). One hemisphere was dissected and stored at −80°C until further processing. The second hemisphere was post-fixed in 4% paraformaldehyde (PFA) in 0.1M phosphate buffer (PB, pH 7.4) for 16 hours then saturated and stored in a cryoprotectant solution (30% sucrose in 0.1% PB, pH 7.4) until sectioned into 40 micrometer thick sections using a freezing sledge microtome (SM 2000R, Leica, Wetzlar, Germany) at − 20°C.

### Immunofluorescence

Briefly, four free floating sections per mouse (n=5 wild-type; n=7 *Mbd2*−/−) were blocked with 10% normal goat serum (NGS) in phosphate buffered saline-0.1% Tween 20 (PBS-T) for 1 hour at room temperature and then incubated with an Alexa Fluor 488-conjugated anti-NeuN antibody (1:200, MAB377X, Millipore, Etobicoke, ON) in 5% NGS in PBS-T overnight at 4°C. For Mbd2 immunostaining the following lentivirus infusion (n=7 scrambled control; n=6 sh-Mbd2), sections were incubated overnight at 4°C with anti-Mbd2 antibody (1:100, Abcam, UK) in 5% NGS in PBS-T, followed by incubation with Alexa Fluor 594-conjugated secondary antibody for 2 hours at room temperature. Sections were then incubated in a Sudan black B staining solution (0.3% Sudan black in 70% EtOH) for 5 minutes. Finally, sections were washed 3 times for 5 minutes in PBS-T and 3 times for 5 minutes in PBS. Sections were then mounted, dried and coverslipped with Aqua Polymount (Polysciences Inc., Warrington, PA).

### Image Analysis

To quantify neuron-number, three regions centered over the appropriate cellular layer CA1, CA2, CA3 were imaged on each of the four sections per animal using an Axio Imager M2 microscope and ZenPro software (Carl Zeiss Canada). Images were saved with 16-bit depth and were processed and analyzed using NIH ImageJ software (National Institutes of Health, USA).

To demonstrate virus targeting and Mbd2 downregulation, 15μm tiled z-stacks of the dorsal hippocampus were acquired with a 20x objective using an LSM710 Confocal Laser Scanning Microscope (Carl Zeiss AG, Germany). Images were then converted to maximum intensity projections using Zen 202 SP5 Black (Carl Zeiss AG, Germany). Acquisition and display settings were held constant for comparative images. Brightness and contrast adjustments were applied to whole images and were held constant across groups.

### Western Blot of Hippocampus tissue

Hippocampi were harvested, lysed in RIPA buffer, loaded into 12% SDS-PAGE gel and transferred to nitrocellulose membrane. Membranes were then probed with antibodies against Drd2 (1:200, Bioss, Woburn, MA), Pogz (1:1000, Bethyl, Montgomery, TX) and Zbtb16 (1:250, Santa-Cruz, Mississauga, ON) Beta-actin was used as a reference protein (1:5000, Sigma-Aldrich, Oakville, ON). Blots were scanned by Amersham Imager 600 imaging system (GE Healthcare, Canada). Possible outliers were visible in the data; therefore, we applied the Iglewicz and Hoaglin’s test for outliers under the strict criterion of z-score ≥|3.5|, which resulted in exclusion of no more than a single data point from each experiment.

### Statistical Analysis

Data are expressed as mean ±SEM unless otherwise stated. Behavioral data were analyzed by two-way ANOVAs with genotype (wild-type/*Mbd2*−/−) and sex as the main factors, followed by HSD Tukey post-hoc tests. Comparisons between two groups were done with two-tailed t-test. qPCR and Bisulfite-Converted PCR-amplicons results for validation of gene-expression and DNA methylation were analyzed as one-tailed tests, based on the a-priori assumption to find differences in the same direction observed by the preceding RNA-seq and capture bisulfite sequencing experiments. In addition to a between-group, a one-sample t-test was applied on data from the object-location memory test to assess if novel-location-memory performance is above chance (50%) within each group. For correlation analysis between DNA methylation and gene expression, the Pearson’s correlation coefficient was calculated (cutoffs: differential methylation |≥25%| and log2 fold-change gene-expression |≥ 0.27|). Other correlations were calculated with Pearson’s R coefficient or Spearman’s Rho coefficient as detailed in the results section. Other statistical analyses of bioinformatic data are detailed in the results section. Heatmaps were generated with Morpheus (https://software.broadinstitute.org/morpheus/; Broad Institute). GO annotations, pathway enrichment analysis and gene-networks were analyzed with Metascape [91] (http://metascape.org).

## Acknowledgements

The work was funded by a grant (PSR-SIIRI) from the Ministère du Développement économique et de l’Innovation, Gouvernement du Québec and a grant from the Canadian Institute for Health Research PJT-159583. EL was funded by the Richard and Edith Strauss Canada Fund post-doctoral fellowship. SDC is the holder of the Charles E. Frosst/Merck Research Associate position. LAW is the recipient of a Doctoral Training Fellowship from the Fonds de recherche du Québec – Santé. The authors wish to thank Dr. David Cheishvili for critical discussion as well as to Bruktawit Maru, Helen Liu, Isabel Kalaycioglu and Stéphanie L’Écuyer for their technical help. Capture and hybridization was performed by the Institut de recherches cliniques de Montréal (Montreal, Canada; licensed by Roche Nimblegen). Next-generation sequencing was performed at Genome-Quebec (Montreal, Canada; licensed by Illumina).

## Supplementary figures legend

**Supplementary Figure 1**. A. Behavioral analyses of male and female *Mbd2*−/− and control mice A. Locomotion in Open-field box was assessed for 5 min. B. Number of entries to center in the open-field test. C. Average speed during the open-field test. D. Social interaction, mice were introduced to a novel mouse for 5 min and interaction time was recorded. E. Exploration time during object-location memory training and F. Discrimination ratio in object-location memory test. G. Time in the light compartment of the Dark-Light Box and H. number of entries to the light side. Numbers within bars represent sample size.

**Supplementary Figure 2**. Analysis of self-grooming behaviors: grooming duration, number of grooming bouts and average bout duration in male and female wild-type and Mbd2−/− mice. A two-way ANOVA found a main effect of sex F(1,28)=7.09, p= 0.0128 on average grooming bout duration. Subsequent independent t-tests found significantly shorter grooming bout duration in wild-type females compared to males.

**Supplementary Figure 3**. Top: Y-maze spontaneous alteration. SAP-Spontaneous Alteration Performance, AAR-Alternate Arm Return, SAR-Same Arm Return, B. Bottom: Exploration is expressed as number of arm entries. Numbers within bars represent sample size.

**Supplementary Figure 4**. Average speed (left), entries to center (right) in the open-field box test. p>0.05 in all cases.

**Supplementary Figure 5**. A. De-novo motif discovery sequences found in HOMER analysis for Mbd2 ChIP-seq peaks. B. Known motif discovery sequences found in HOMER analysis for Mbd2 ChIP-seq peaks

**Supplementary Figure 6**. A. GO-Pathway analysis enrichment of Mbd2-bound genes. B. Gene-network analysis of Mbd2 binding peaks.

**Supplementary Figure 7**. A. Q-ChIP validation of Mbd2 peaks. B. Q-ChIP for RNApolII (SP5) and C. H3K4me1. For Q-ChIP of Mbd2, RNApolII(SP5) and H3K4me1 pool of 5-6 mice analyzed by triplicate/group). Data are presented as mean ±SEM # p<0.1, *p<0.05, **p<0.01 (T-test). Dashed lines represent Mbd2−/− binding levels.

**Supplementary Figure 8**. A. A capture from genome browser showing RNA-seq read alignments for the first 3 exons of the *Mbd2* gene. *Mbd2*−/− mice show expression of the first exon only, as expected by the design of this mouse line [24]. B. QPCR validation for down-regulated genes (*Apba2, Kcnd2, Clstn1, Gria3a, Jarid2, Gria1*) and up-regulated genes (*Ttr, Aqp1, Rdh5*) shows change in gene-expression in *Mbd2*−/− mice in agreement with the RNA-Seq data (n=9 wild-type, n=6 *Mbd2*−/−; *p<0.05, **p<0.01 t-test).

**Supplementary Figure 9**. A. GO-Pathway analysis enrichment of *Mbd2*−/− down-regulated genes. B. Gene-network analysis *Mbd2*−/− down-regulated genes.

**Supplementary Figure 10**. A. GO-Pathway analysis enrichment of *Mbd2*−/− up-regulated genes. B. Gene-network analysis *Mbd2*−/− up-regulated genes.

**Supplementary Figure 11**. Effect of *Mbd2*−/− on gene expression of epigenetic readers and modifiers. *FDR-corrected p<0.05, **FDR-corrected p<0.01, ^#^FDR-corrected p<0.1. Note: Mbd2-aligned reads in *Mbd2*−/− mice are from the intact exon 1 (see also Fig S6A).

**Supplementary Figure 12**. A. A table summarizing the number of differentially methylated CpGs and genes under the different analytical conditions. B. A capture from IGV genomic browser depicting Mbd2 binding peak around Sfi1 transcription start site (TSS) with methylation levels and differential-methylation between wild-type and *Mbd2*−/− mice hippocampus. C. cumulative distribution of methylation levels of regulatory DNA regions genome-wide demonstrating hyper-methylation in *Mbd2*−/− hippocampus (K-S test). D. Significant motifs discovered for promoter CpG with positive correlation between DNA methylation and gene-expression are depicted. No significant motifs were found for CpGs with negative correlation between DNA methylation and gene-expression. E-F. Pathway analysis of hypo-(E) and hyper-(F) methylated gene promoters.

**Supplementary Figure 13**. A. Correlation analysis of CpG methylation levels in capture-array genome-wide bisulfite-sequencing and amplicons bisulfite-sequencing for wild-type mice. B. Correlation analysis of CpG methylation levels in capture-array genome-wide bisulfite-sequencing and amplicons bisulfite-sequencing for wild-type mice. C. Average change in methylation levels across all CpGs (with at least 1% change in methylation) show larger change in methylation between Mbd2−/− and wild-type mice in CpGs that became hyper-methylated (n=114) than in CpGs that became hypo-methylated (n=80) (t-test, **p<0.01) D. More CpGs with at least 1% change in methylation between mbd2−/− and wild-type mice are hyper-methylated than hypo-methylated (binomial test, *p<0.05). E. A cumulative distribution of methylation levels within Mbd2 peak regions in a validation cohort (K-S test).

**Supplementary Figure 14**. A-B. Protein-Protein Interaction Networks for down-(A) and up-(B) regulated genes overlapped with ASD risk genes (Sanders et al[52]). PPI=Protein-Protein Interaction.

**Supplementary Figure 15**. A-B. Protein-Protein Interaction Networks for down-(A) and up-(B) regulated genes in *Mbd2*−/− overlapped with SFARI-GENE dataset. PPI=Protein-Protein Interaction. Disconnected nodes are omitted from (A) for clarity.

**Supplementary Figure 16**. Correlation analyses of neuron count and neuron soma size (top); neuron count and layer thickness (middle) and neuron soma size and layer thickness (bottom) of hippocampal subregions CA1-CA3 in wild-type and *Mbd2*−/− mice as determined by quantitative analyses following NeuN immunolabeling. Correlation analyses between neuron count and neuron soma size (left);neuron count and layer thickness (middle) and between neuron soma size and layer thickness (right) of the total hippocampus (collapsing CA1, CA2 and CA3 data). Spearman’s Rho for correlation analyses, with exact p-values indicated in each graph.

**Supplementary Figure 17**. A. Schematic illustration of the sh-*Mbd2*/scramble constructs. RRE-Rev Response Element, hCMV-human Cytomegalovirus promoter, tGFP-turbo GFP reporter, IRES-Internal Ribosomal Entry Site, PURO^R^-Puromycin Resistance, WPRE-Woodchuck Hepatitis Posttranslational Regulatory Element. B. In-vitro validation of sh-*Mbd2* construct. The effect of sh-*Mbd2* constructs on Mbd2 mRNA levels in lentivirus-treated NIH3T3 was tested by qRT-PCR (change over scramble control). C. Illustration of infusion site adapted from the Paxinos and Franklin mouse brain atlas (top) and confocal fluorescence microscope images showing hippocampus expression of GFP-tagged lentivirus (green), MBD2 (red) and a merged image (center), scale bar: 200μm; and larger magnification of lentivirus-infected neurons (bottom), scale bar: 50 μm.

## References

1. Sweatt, J.D., Neural plasticity and behavior - sixty years of conceptual advances. J Neurochem, 2016. 139 Suppl 2: p. 179–199.

2. Lister, R., et al., Human DNA methylomes at base resolution show widespread epigenomic differences. Nature, 2009. 462(7271): p. 315–22.

3. Weber, M., et al., Distribution, silencing potential and evolutionary impact of promoter DNA methylation in the human genome. Nat Genet, 2007. 39(4): p. 457–66.

4. Du, Q., et al., Methyl-CpG-binding domain proteins: readers of the epigenome. Epigenomics, 2015. 7(6): p. 1051–73.

5. Jorgensen, H.F. and A. Bird, MeCP2 and other methyl-CpG binding proteins. Ment Retard Dev Disabil Res Rev, 2002. 8(2): p. 87–93.

6. Baubec, T., et al., Methylation-dependent and - independent genomic targeting principles of the MBD protein family. Cell, 2013. 153(2): p. 480–92.

7. Nan, X., F.J. Campoy, and A. Bird, MeCP2 is a transcriptional repressor with abundant binding sites in genomic chromatin. Cell, 1997. 88(4): p. 471–81.

8. Ng, H.H., et al., MBD2 is a transcriptional repressor belonging to the MeCP1 histone deacetylase complex. Nat Genet, 1999. 23(1): p. 58–61.

9. Fan, G. and L. Hutnick, Methyl-CpG binding proteins in the nervous system. Cell Res, 2005. 15(4): p. 255–61.

10. Hendrich, B. and A. Bird, Identification and characterization of a family of mammalian methyl-CpG binding proteins. Mol Cell Biol, 1998. 18(11): p. 6538–47.

11. Amir, R.E., et al., Rett syndrome is caused by mutations in X-linked MECP2, encoding methyl-CpG-binding protein 2. Nat Genet, 1999. 23(2): p. 185–8.

12. Moretti, P. and H.Y. Zoghbi, MeCP2 dysfunction in Rett syndrome and related disorders. Curr Opin Genet Dev, 2006. 16(3): p. 276–81.

13. Zhao, X., et al., Mice lacking methyl-CpG binding protein 1 have deficits in adult neurogenesis and hippocampal function. Proc Natl Acad Sci U S A, 2003. 100(11): p. 6777–82.

14. Dong, E., et al., DNA-methyltransferase1 (DNMT1) binding to CpG rich GABAergic and BDNF promoters is increased in the brain of schizophrenia and bipolar disorder patients. Schizophr Res, 2015. 167(1-3): p. 35–41.

15. Gandal, M.J., et al., Shared molecular neuropathology across major psychiatric disorders parallels polygenic overlap. Science, 2018. 359(6376): p. 693–697.

16. Li, H., et al., Mutation analysis of methyl-CpG binding protein family genes in autistic patients. Brain Dev, 2005. 27(5): p. 321–5.

17. Coe, B.P., et al., Neurodevelopmental disease genes implicated by de novo mutation and copy number variation morbidity. Nat Genet, 2019. 51(1): p. 106–116.

18. Nagy, C., et al., Astrocytic abnormalities and global DNA methylation patterns in depression and suicide. Mol Psychiatry, 2015. 20(3): p. 320–8.

19. Weaver, I.C., et al., The methylated-DNA binding protein MBD2 enhances NGFI-A (egr-1)- mediated transcriptional activation of the glucocorticoid receptor. Philos Trans R Soc Lond B Biol Sci, 2014. 369(1652).

20. Moretti, P., et al., Learning and memory and synaptic plasticity are impaired in a mouse model of Rett syndrome. J Neurosci, 2006. 26(1): p. 319–27.

21. Li, H., et al., Cell cycle-linked MeCP2 phosphorylation modulates adult neurogenesis involving the Notch signalling pathway. Nat Commun, 2014. 5: p. 5601.

22. Guy, J., et al., Reversal of neurological defects in a mouse model of Rett syndrome. Science, 2007. 315(5815): p. 1143–7.

23. Jobe, E.M., et al., Methyl-CpG-Binding Protein MBD1 Regulates Neuronal Lineage Commitment through Maintaining Adult Neural Stem Cell Identity. J Neurosci, 2017. 37(3): p. 523–536.

24. Hendrich, B., et al., Closely related proteins MBD2 and MBD3 play distinctive but interacting roles in mouse development. Genes Dev, 2001. 15(6): p. 710–23.

25. Wood, K.H., et al., Tagging methyl-CpG-binding domain proteins reveals different spatiotemporal expression and supports distinct functions. Epigenomics, 2016. 8(4): p. 455–73.

26. McNaughton, C.H., et al., Evidence for social anxiety and impaired social cognition in a mouse model of fragile X syndrome. Behav Neurosci, 2008. 122(2): p. 293–300.

27. Mineur, Y.S., L.X. Huynh, and W.E. Crusio, Social behavior deficits in the Fmr1 mutant mouse. Behav Brain Res, 2006. 168(1): p. 172–5.

28. Sato, A., et al., Rapamycin reverses impaired social interaction in mouse models of tuberous sclerosis complex. Nat Commun, 2012. 3: p. 1292.

29. Tavares, R.M., et al., A Map for Social Navigation in the Human Brain. Neuron, 2015. 87(1): p. 231–43.

30. Yang, L., et al., Hypocretin/orexin neurons contribute to hippocampus-dependent social memory and synaptic plasticity in mice. J Neurosci, 2013. 33(12): p. 5275–84.

31. Heyward, F.D., et al., Obesity Weighs down Memory through a Mechanism Involving the Neuroepigenetic Dysregulation of Sirt1. J Neurosci, 2016. 36(4): p. 1324–35.

32. Heyward, F.D., et al., Adult mice maintained on a high-fat diet exhibit object location memory deficits and reduced hippocampal SIRT1 gene expression. Neurobiol Learn Mem, 2012. 98(1): p. 25–32.

33. Mineur, Y.S., et al., Cholinergic signaling in the hippocampus regulates social stress resilience and anxiety- and depression-like behavior. Proc Natl Acad Sci U S A, 2013. 110(9): p. 3573–8.

34. Camarena, V., et al., Disruption of Mbd5 in mice causes neuronal functional deficits and neurobehavioral abnormalities consistent with 2q23.1 microdeletion syndrome. EMBO Mol Med, 2014. 6(8): p. 1003–15.

35. Lu, H., et al., Loss and Gain of MeCP2 Cause Similar Hippocampal Circuit Dysfunction that Is Rescued by Deep Brain Stimulation in a Rett Syndrome Mouse Model. Neuron, 2016. 91(4): p. 739–747.

36. de Bruin, A., et al., Genome-wide analysis reveals NRP1 as a direct HIF1alpha-E2F7 target in the regulation of motorneuron guidance in vivo. Nucleic Acids Res, 2016. 44(8): p. 3549–66.

37. Ghanem, N., et al., The Rb/E2F pathway modulates neurogenesis through direct regulation of the Dlx1/Dlx2 bigene cluster. J Neurosci, 2012. 32(24): p. 8219–30.

38. Serrano-Perez, M.C., et al., NFAT transcription factors regulate survival, proliferation, migration, and differentiation of neural precursor cells. Glia, 2015. 63(6): p. 987–1004.

39. Quadrato, G., et al., Modulation of GABAA receptor signaling increases neurogenesis and suppresses anxiety through NFATc4. J Neurosci, 2014. 34(25): p. 8630–45.

40. Gjoneska, E., et al., Conserved epigenomic signals in mice and humans reveal immune basis of Alzheimer’s disease. Nature, 2015. 518(7539): p. 365–9.

41. Hirose, Y. and Y. Ohkuma, Phosphorylation of the C-terminal domain of RNA polymerase II plays central roles in the integrated events of eucaryotic gene expression. J Biochem, 2007. 141(5): p. 601–8.

42. Jonkers, I. and J.T. Lis, Getting up to speed with transcription elongation by RNA polymerase II. Nat Rev Mol Cell Biol, 2015. 16(3): p. 167–77.

43. Creyghton, M.P., et al., Histone H3K27ac separates active from poised enhancers and predicts developmental state. Proc Natl Acad Sci U S A, 2010. 107(50): p. 21931–6.

44. Heintzman, N.D., et al., Histone modifications at human enhancers reflect global cell-type-specific gene expression. Nature, 2009. 459(7243): p. 108–12.

45. Pundhir, S., et al., Peak-valley-peak pattern of histone modifications delineates active regulatory elements and their directionality. Nucleic Acids Res, 2016. 44(9): p. 4037–51.

46. McGhee, J.D., et al., A 200 base pair region at the 5’ end of the chicken adult beta-globin gene is accessible to nuclease digestion. Cell, 1981. 27(1 Pt 2): p. 45–55.

47. Ludwig, A.K., P. Zhang, and M.C. Cardoso, Modifiers and Readers of DNA Modifications and Their Impact on Genome Structure, Expression, and Stability in Disease. Front Genet, 2016. 7: p. 115.

48. Wang, L., et al., Mbd2 promotes foxp3 demethylation and T-regulatory-cell function. Mol Cell Biol, 2013. 33(20): p. 4106–15.

49. Yin, Y., et al., Impact of cytosine methylation on DNA binding specificities of human transcription factors. Science, 2017. 356(6337).

50. Cukier, H.N., et al., The expanding role of MBD genes in autism: identification of a MECP2 duplication and novel alterations in MBD5, MBD6, and SETDB1. Autism Res, 2012. 5(6): p. 385–97.

51. Gandal, M.J., et al., Transcriptome-wide isoform-level dysregulation in ASD, schizophrenia, and bipolar disorder. Science, 2018. 362(6420).

52. Sanders, S.J., et al., Insights into Autism Spectrum Disorder Genomic Architecture and Biology from 71 Risk Loci. Neuron, 2015. 87(6): p. 1215–1233.

53. Cheishvili, D., et al., Synergistic effects of combined DNA methyltransferase inhibition and MBD2 depletion on breast cancer cells; MBD2 depletion blocks 5-aza-2’-deoxycytidine-triggered invasiveness. Carcinogenesis, 2014. 35(11): p. 2436–46.

54. Shukeir, N., et al., Alteration of the methylation status of tumor-promoting genes decreases prostate cancer cell invasiveness and tumorigenesis in vitro and in vivo. Cancer Res, 2006. 66(18): p. 9202–10.

55. Stefanska, B., et al., Transcription onset of genes critical in liver carcinogenesis is epigenetically regulated by methylated DNA-binding protein MBD2. Carcinogenesis, 2013. 34(12): p. 2738–49.

56. Edalatmanesh, M.A., et al., Increased hippocampal cell density and enhanced spatial memory in the valproic acid rat model of autism. Brain Res, 2013. 1526: p. 15–25.

57. Sosa-Diaz, N., et al., Prefrontal cortex, hippocampus, and basolateral amygdala plasticity in a rat model of autism spectrum. Synapse, 2014. 68(10): p. 468–73.

58. Nicolson, R., et al., Detection and mapping of hippocampal abnormalities in autism. Psychiatry Res, 2006. 148(1): p. 11–21.

59. Courchesne, E., et al., Neuron number and size in prefrontal cortex of children with autism. JAMA, 2011. 306(18): p. 2001–10.

60. Berger, J. and A. Bird, Role of MBD2 in gene regulation and tumorigenesis. Biochem Soc Trans, 2005. 33(Pt 6): p. 1537–40.

61. Nan, X., S. Cross, and A. Bird, Gene silencing by methyl-CpG-binding proteins. Novartis Found Symp, 1998. 214: p. 6–16; discussion 16–21, 46–50.

62. Chahrour, M., et al., MeCP2, a key contributor to neurological disease, activates and represses transcription. Science, 2008. 320(5880): p. 1224–9.

63. Li, C., et al., MeCP2 enforces Foxp3 expression to promote regulatory T cells’ resilience to inflammation. Proc Natl Acad Sci U S A, 2014. 111(27): p. E2807–16.

64. Angrisano, T., et al., TACC3 mediates the association of MBD2 with histone acetyltransferases and relieves transcriptional repression of methylated promoters. Nucleic Acids Res, 2006. 34(1): p. 364–72.

65. Detich, N., J. Theberge, and M. Szyf, Promoter-specific activation and demethylation by MBD2/demethylase. J Biol Chem, 2002. 277(39): p. 35791–4.

66. Cui, Y. and J. Irudayaraj, Dissecting the behavior and function of MBD3 in DNA methylation homeostasis by single-molecule spectroscopy and microscopy. Nucleic Acids Res, 2015. 43(6): p. 3046–55.

67. Vogel Ciernia, A. and J. LaSalle, The landscape of DNA methylation amid a perfect storm of autism aetiologies. Nat Rev Neurosci, 2016. 17(7): p. 411–23.

68. Nardone, S., et al., Dysregulation of Cortical Neuron DNA Methylation Profile in Autism Spectrum Disorder. Cereb Cortex, 2017. 27(12): p. 5739–5754.

69. Wong, C.C.Y., et al., Genome-wide DNA methylation profiling identifies convergent molecular signatures associated with idiopathic and syndromic autism in post-mortem human brain tissue. Hum Mol Genet, 2019. 28(13): p. 2201–2211.

70. Ramaswami, G., et al., Integrative genomics identifies a convergent molecular subtype that links epigenomic with transcriptomic differences in autism. Nat Commun, 2020. 11(1): p. 4873.

71. Hannon, E., et al., Elevated polygenic burden for autism is associated with differential DNA methylation at birth. Genome Med, 2018. 10(1): p. 19.

72. Nardone, S. and E. Elliott, The Interaction between the Immune System and Epigenetics in the Etiology of Autism Spectrum Disorders. Front Neurosci, 2016. 10: p. 329.

73. Ladd-Acosta, C., et al., Common DNA methylation alterations in multiple brain regions in autism. Mol Psychiatry, 2014. 19(8): p. 862–71.

74. Richetto, J., et al., Genome-wide DNA Methylation Changes in a Mouse Model of Infection-Mediated Neurodevelopmental Disorders. Biol Psychiatry, 2017. 81(3): p. 265–276.

75. Basil, P., et al., Prenatal immune activation alters the adult neural epigenome but can be partly stabilised by a n-3 polyunsaturated fatty acid diet. Transl Psychiatry, 2018. 8(1): p. 125.

76. Xie, B., et al., Genetic association study between methyl-CpG-binding domain genes and schizophrenia among Chinese family trios. Psychiatr Genet, 2014. 24(5): p. 221–4.

77. Sams, D.S., et al., Neuronal CTCF Is Necessary for Basal and Experience-Dependent Gene Regulation, Memory Formation, and Genomic Structure of BDNF and Arc. Cell Rep, 2016. 17(9): p. 2418–2430.

78. Langmead, B., Aligning short sequencing reads with Bowtie. Curr Protoc Bioinformatics, 2010. Chapter 11: p. Unit 11 7.

79. Liu, T., Use model-based Analysis of ChIP-Seq (MACS) to analyze short reads generated by sequencing protein-DNA interactions in embryonic stem cells. Methods Mol Biol, 2014. 1150: p. 81–95.

80. Tarbell, E.D. and T. Liu, HMMRATAC: a Hidden Markov ModeleR for ATAC-seq. Nucleic Acids Res, 2019. 47(16): p. e91.

81. Dobin, A., et al., STAR: ultrafast universal RNA-seq aligner. Bioinformatics, 2013. 29(1): p. 15–21.

82. Love, M.I., W. Huber, and S. Anders, Moderated estimation of fold change and dispersion for RNA-seq data with DESeq2. Genome Biol, 2014. 15(12): p. 550.

83. Topham, L., et al., The transition from acute to chronic pain: dynamic epigenetic reprogramming of the mouse prefrontal cortex up to 1 year after nerve injury. Pain, 2020. 161(10): p. 2394–2409.

84. Schmidt, M., et al., Fetal glucocorticoid receptor (Nr3c1) deficiency alters the landscape of DNA methylation of murine placenta in a sex-dependent manner and is associated to anxiety-like behavior in adulthood. Transl Psychiatry, 2019. 9(1): p. 23.

85. Xi, Y. and W. Li, BSMAP: whole genome bisulfite sequence MAPping program. BMC Bioinformatics, 2009. 10: p. 232.

86. Akalin, A., et al., methylKit: a comprehensive R package for the analysis of genome-wide DNA methylation profiles. Genome Biol, 2012. 13(10): p. R87.

87. Heinz, S., et al., Simple combinations of lineage-determining transcription factors prime cis-regulatory elements required for macrophage and B cell identities. Mol Cell, 2010. 38(4): p. 576–89.

88. Thomas-Chollier, M., et al., A complete workflow for the analysis of full-size ChIP-seq (and similar) data sets using peak-motifs. Nat Protoc, 2012. 7(8): p. 1551–68.

89. Lu, J., et al., PrimerSuite: A High-Throughput Web-Based Primer Design Program for Multiplex Bisulfite PCR. Sci Rep, 2017. 7: p. 41328.

90. Szklarczyk, D., et al., The STRING database in 2017: quality-controlled protein-protein association networks, made broadly accessible. Nucleic Acids Res, 2017. 45(D1): p. D362–D368.

91. Tripathi, S., et al., Meta- and Orthogonal Integration of Influenza “OMICs” Data Defines a Role for UBR4 in Virus Budding. Cell Host Microbe, 2015. 18(6): p. 723–35.

