## Supplementary figures and images for "Methyl-CpG binding domain 2 (Mbd2) is an Epigenetic Regulator of Autism-Risk Genes and Cognition"

### Supplemental Figure 1

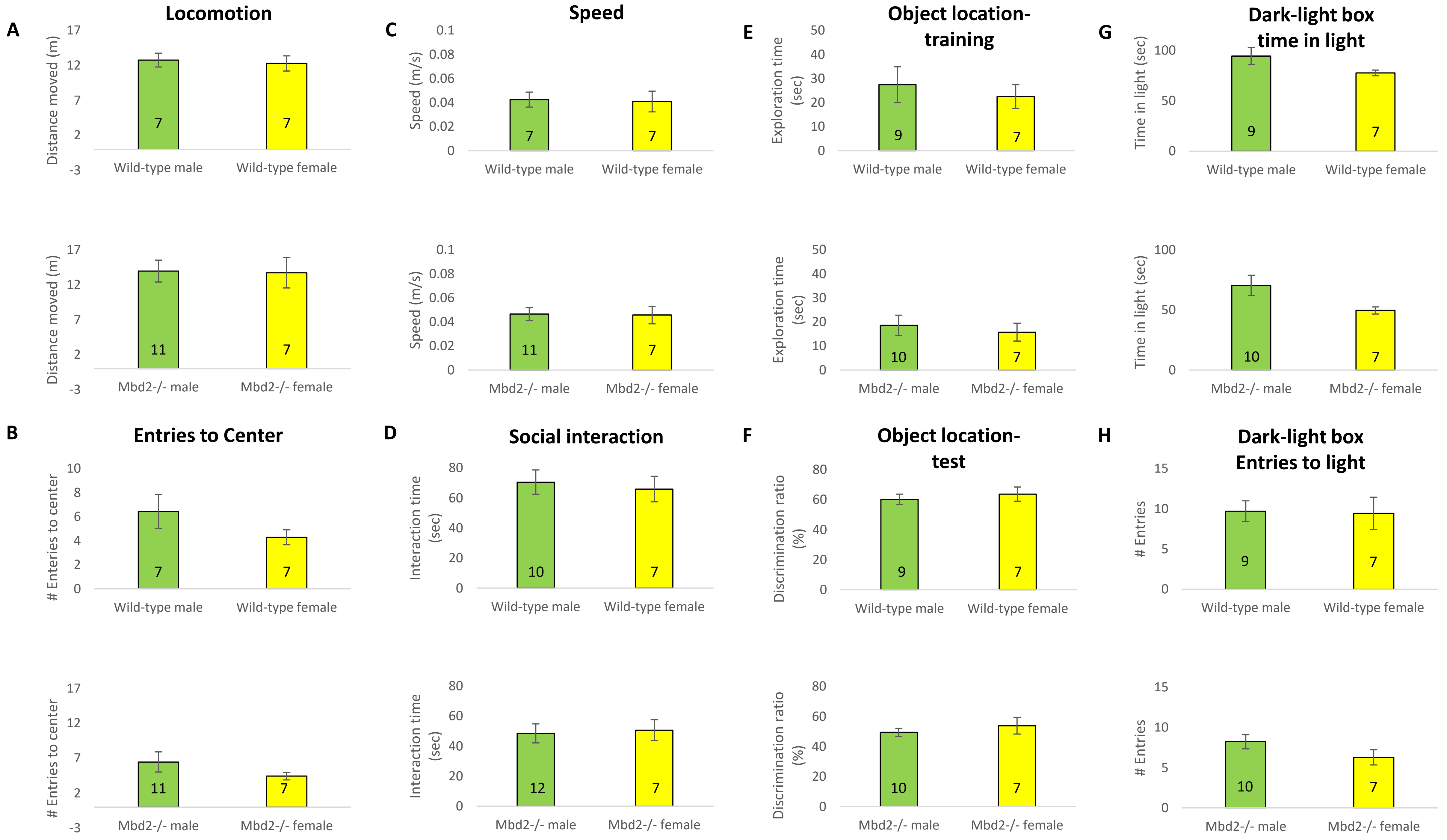

### Supplemental Figure 2

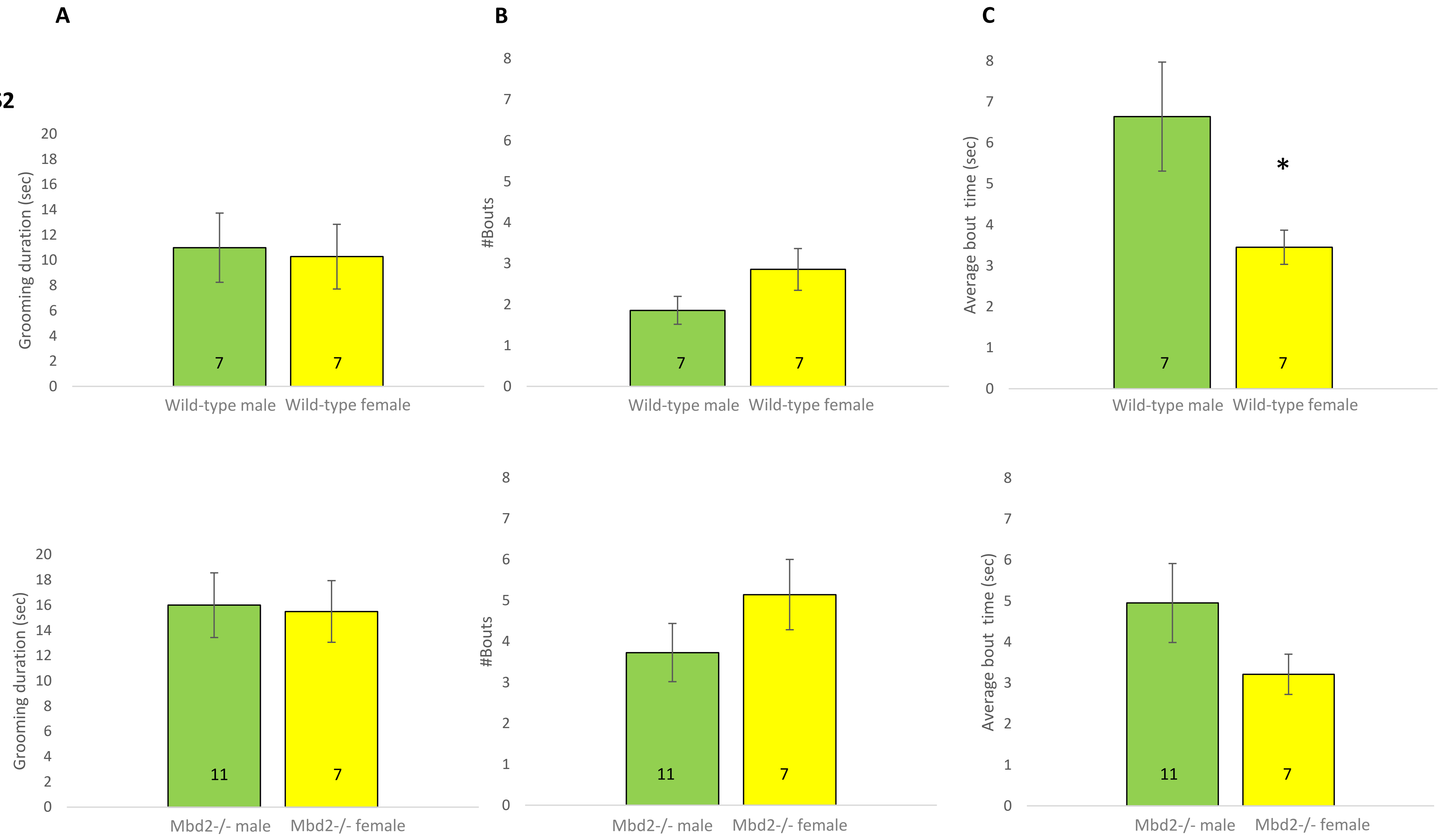

### Supplemental Figure 3

**Fig.S3**

#### Y Maze Spontaneous Alternation Test

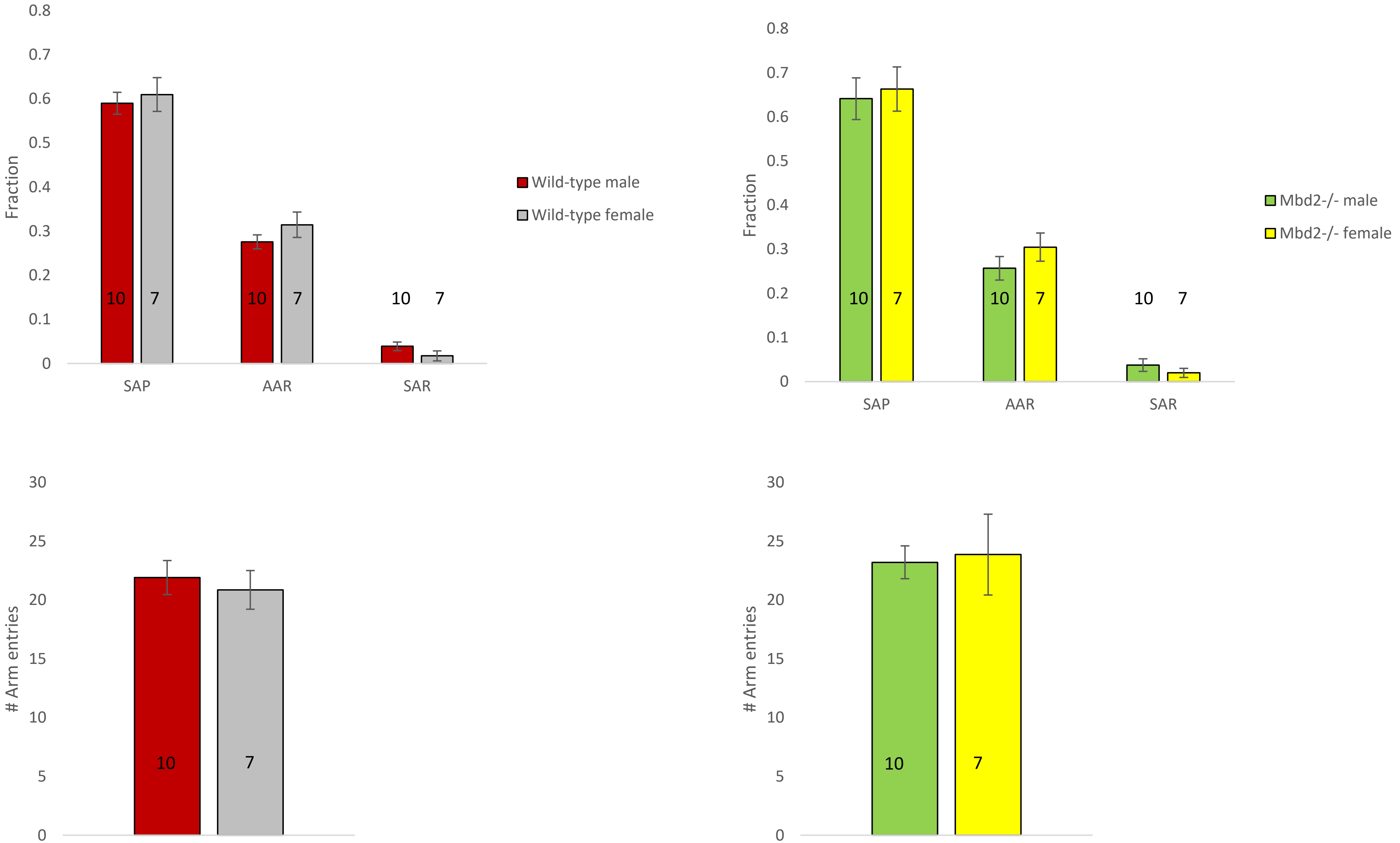

### Supplemental Figure 4

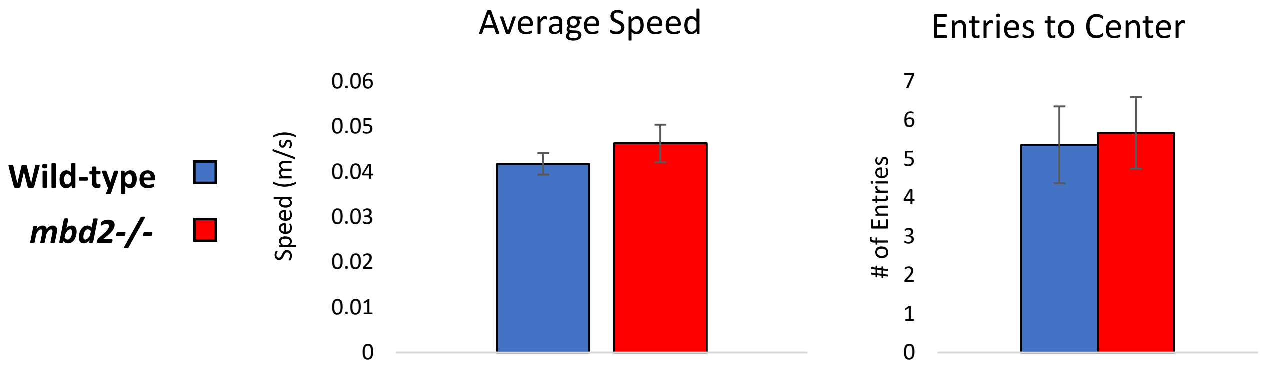

### Supplemental Figure 6

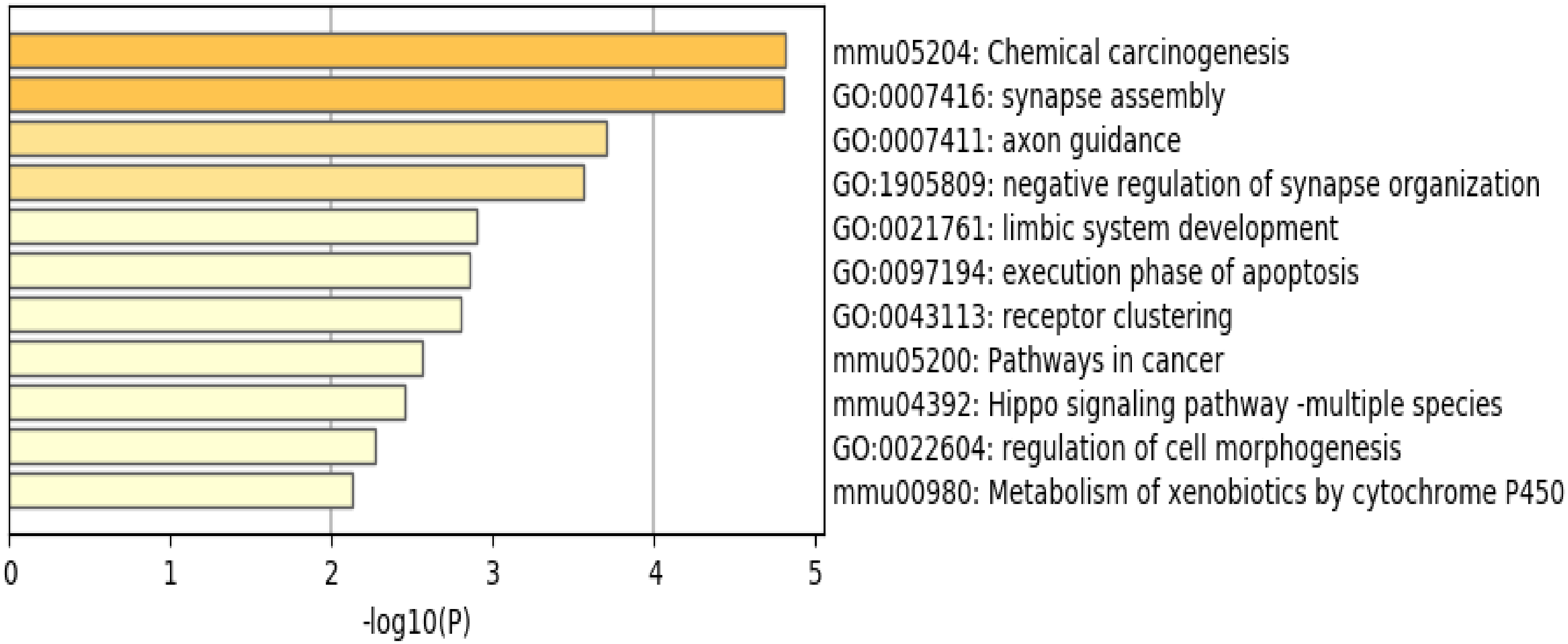

### Supplemental Figure 7

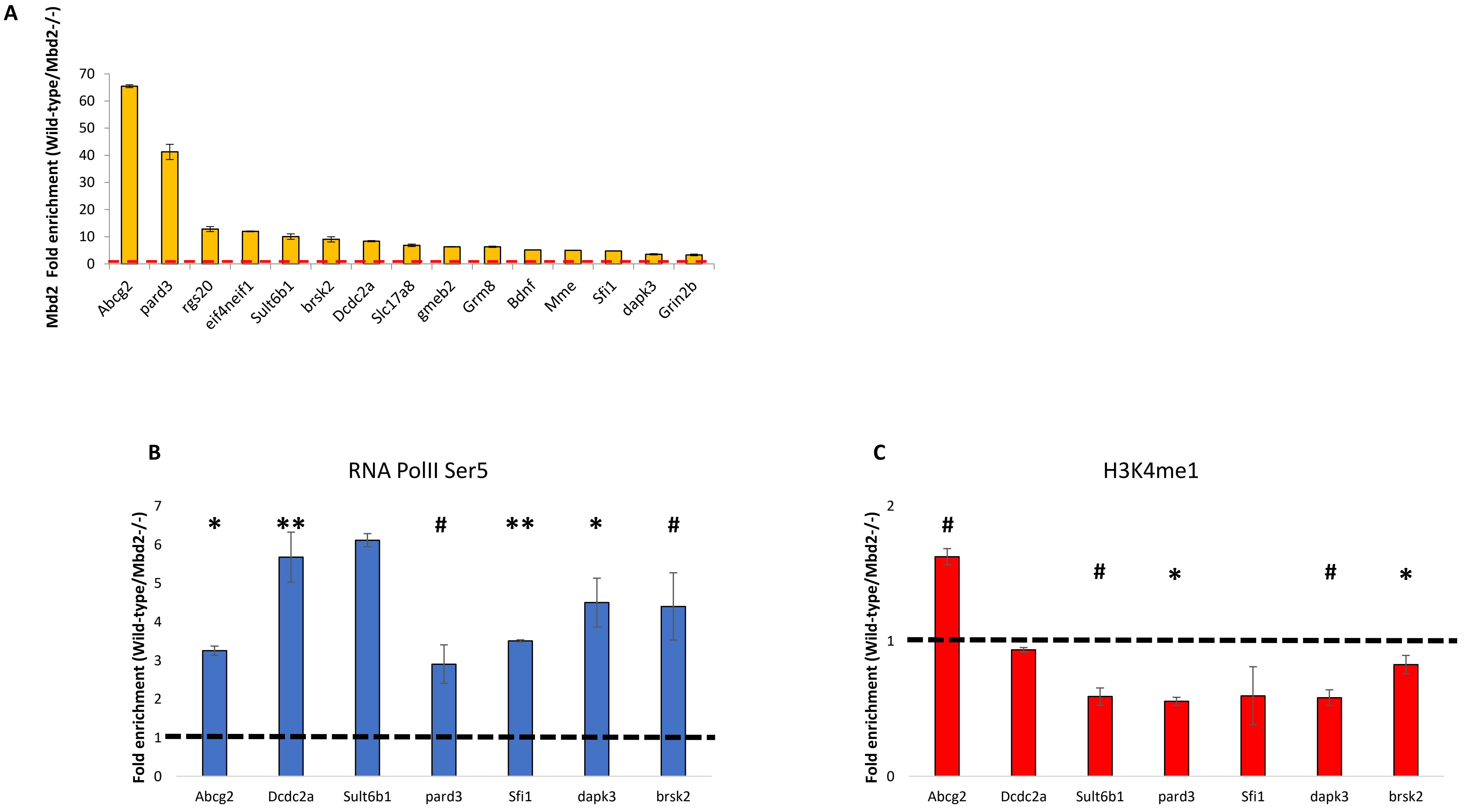

### Supplemental Figure 8

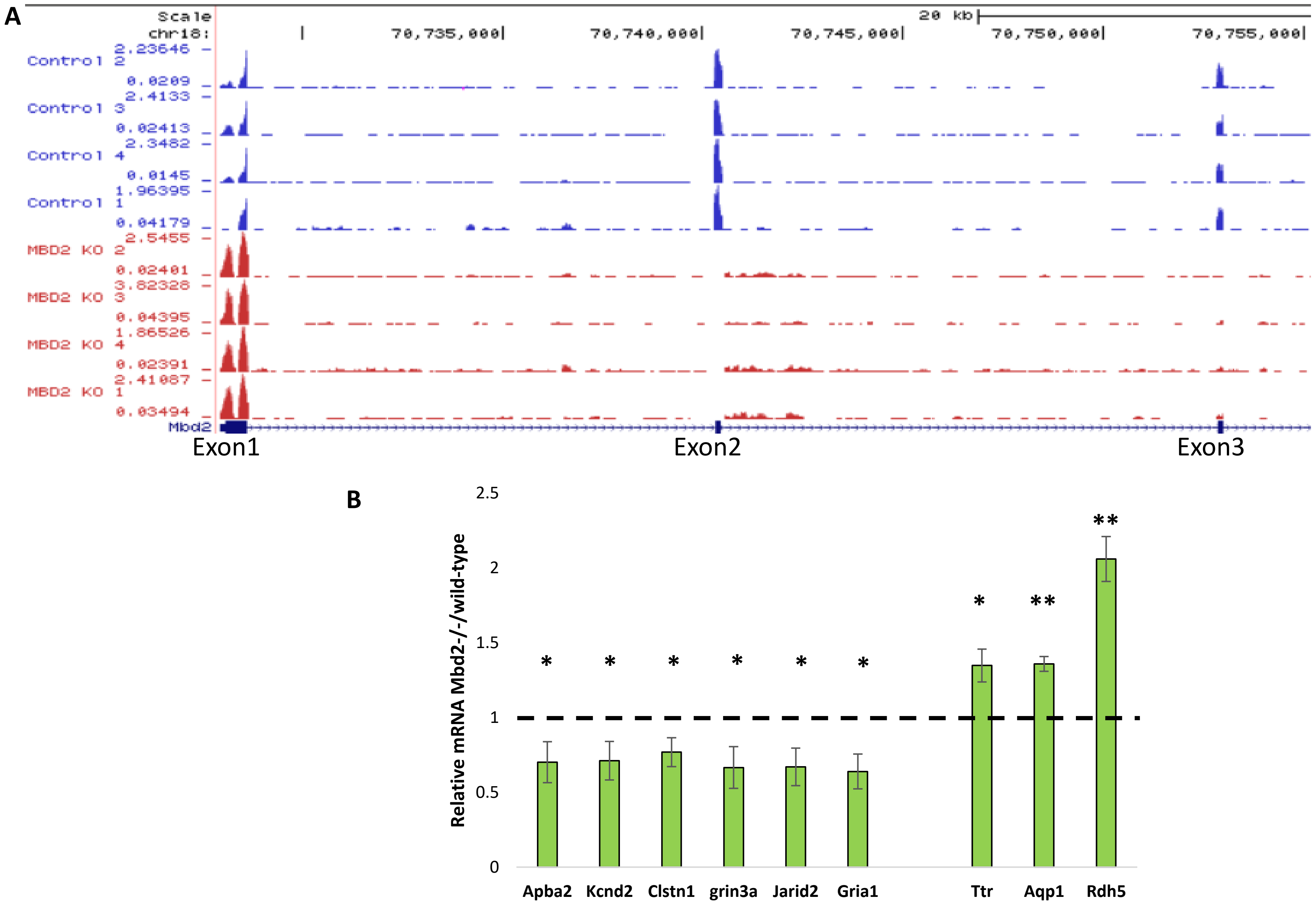

### Supplemental Figure 9

**Fig S9** **Down-regulated genes**

**A**

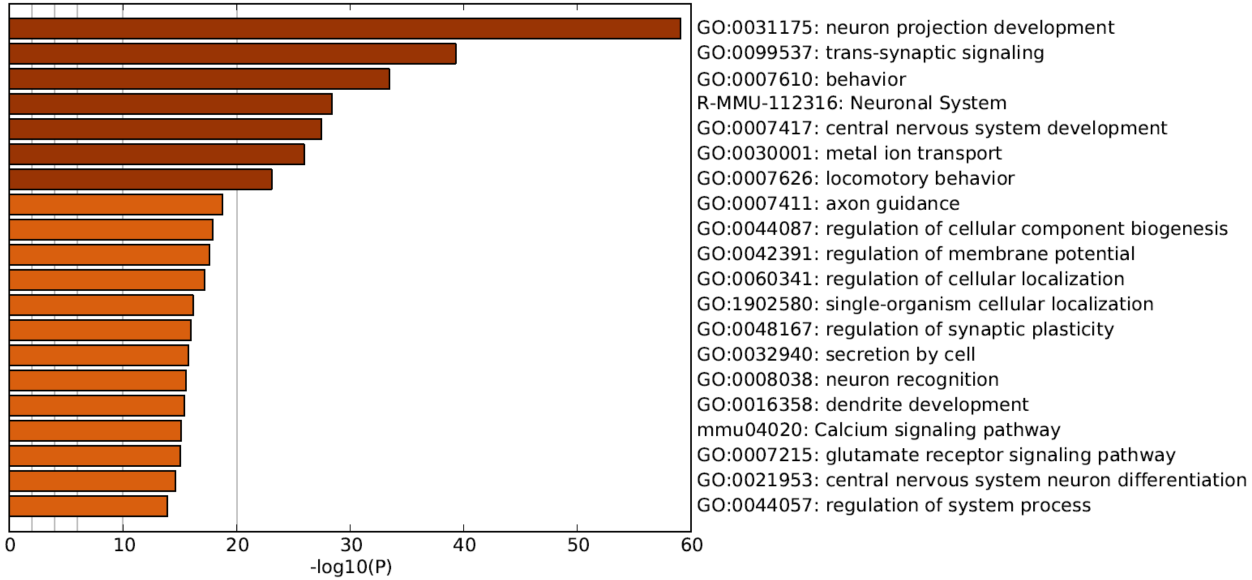


**B**

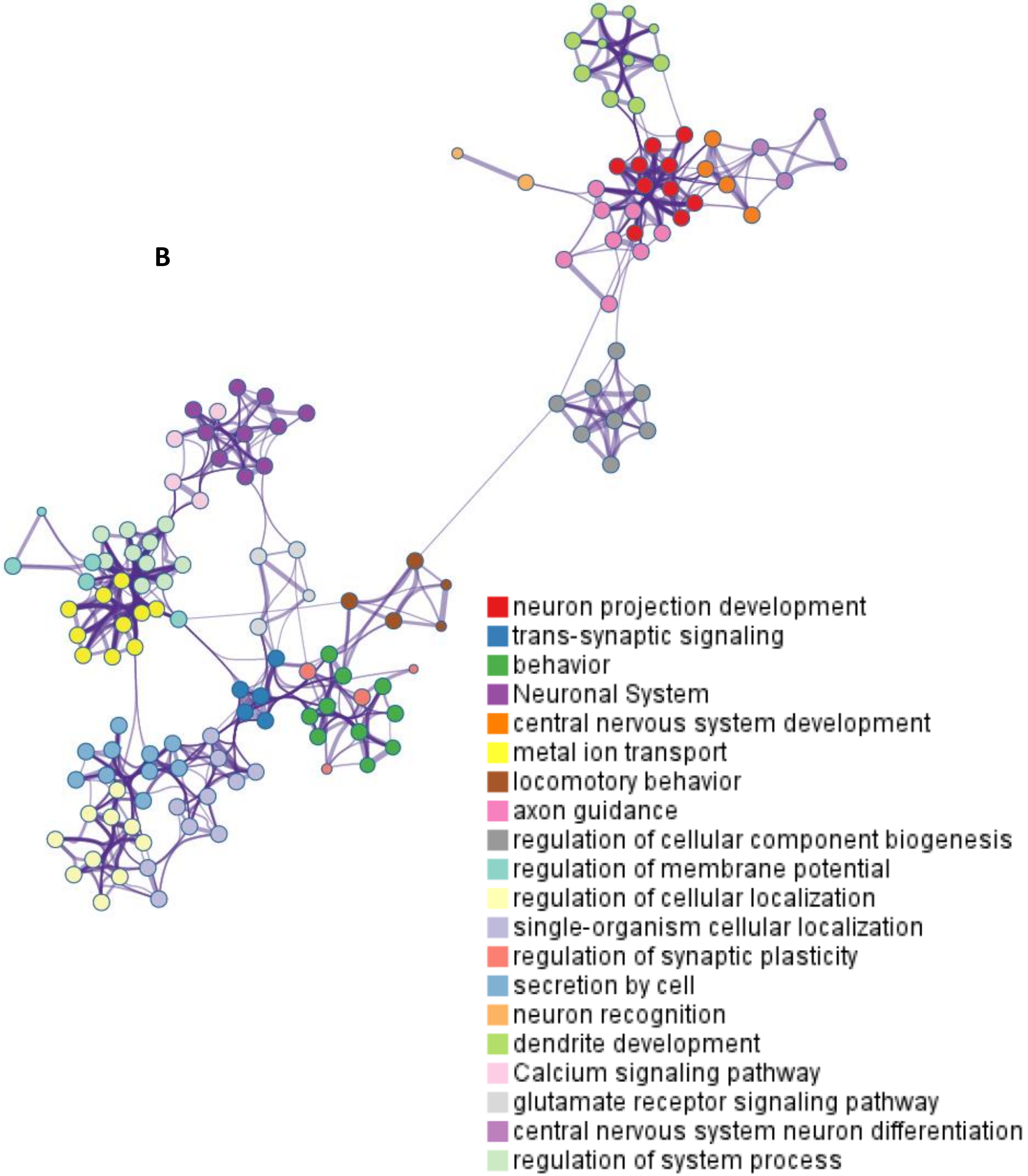

### Supplemental Figure 10

**Fig S10** **Up-regulated genes**

**B**

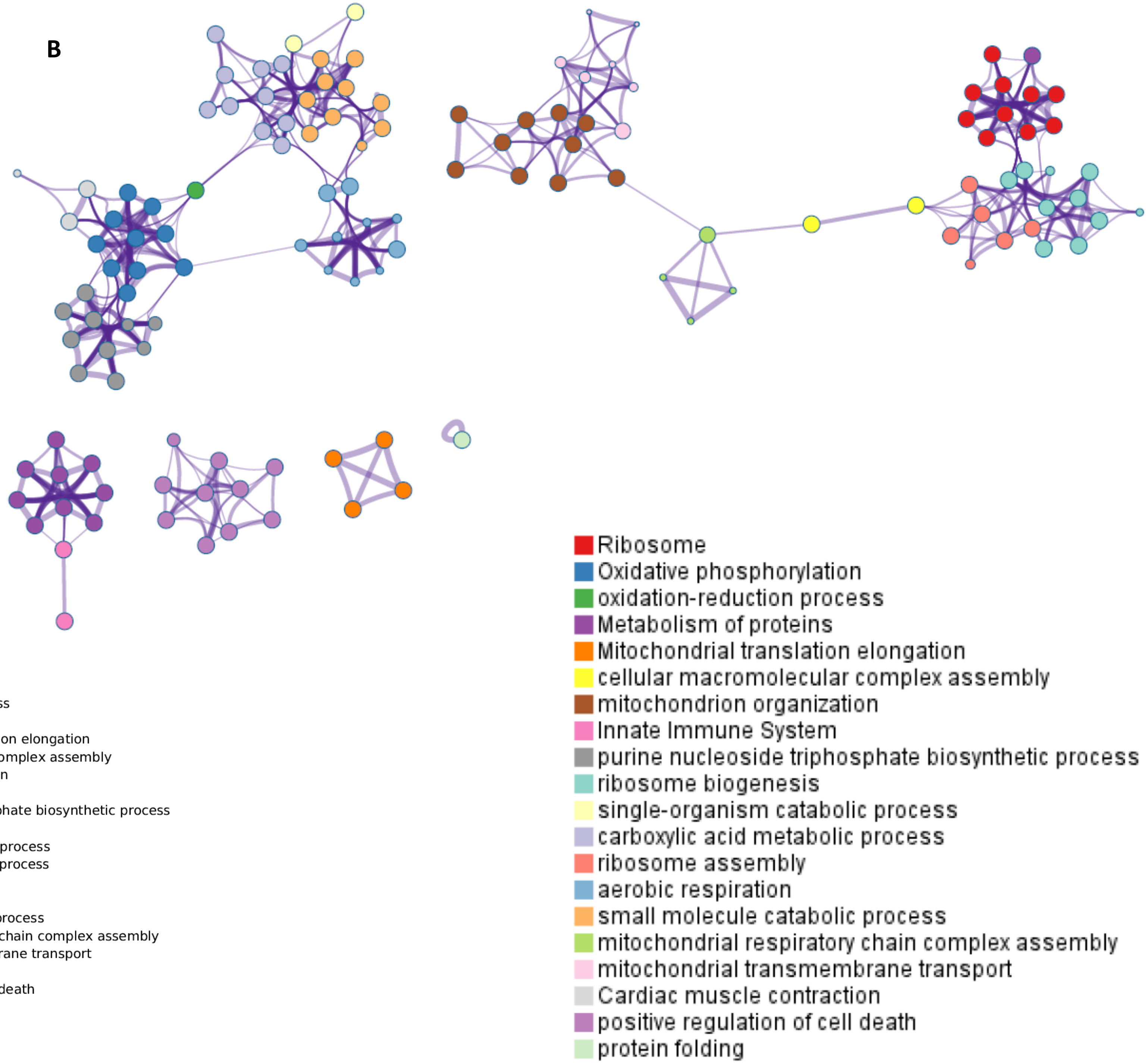


**A**

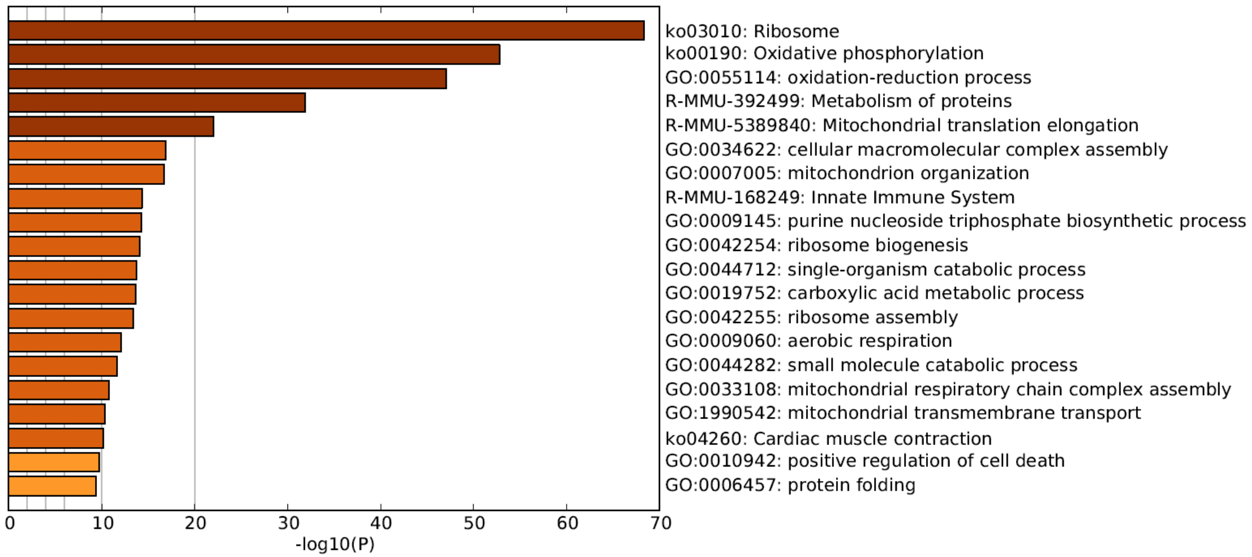

### Supplemental Figure 11

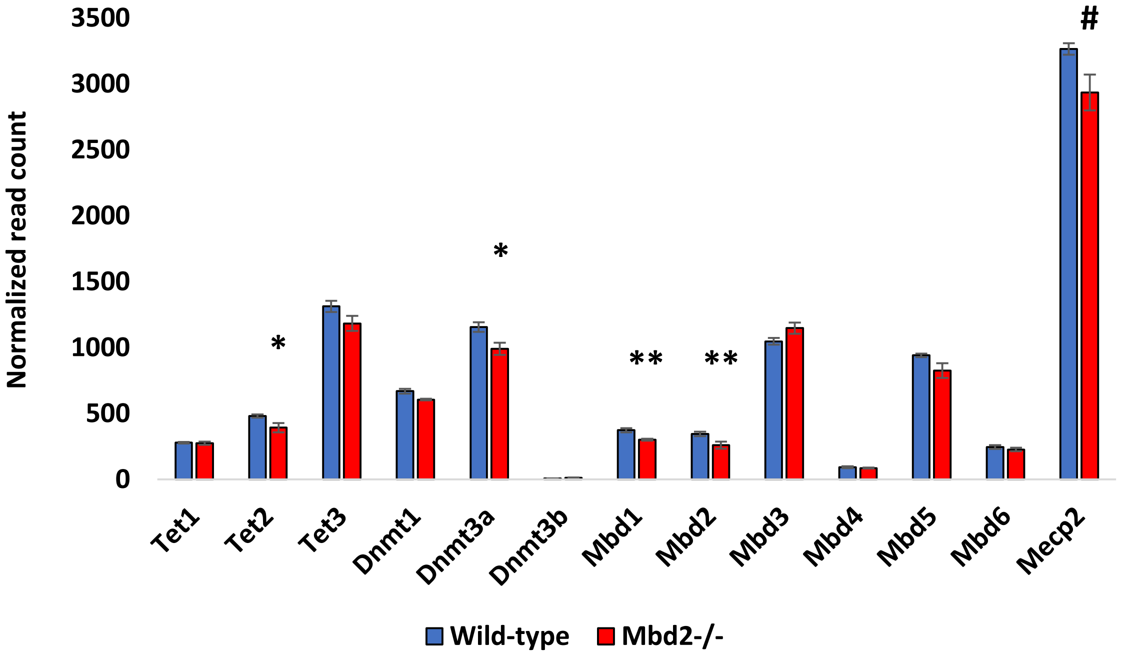

### Supplemental Figure 13

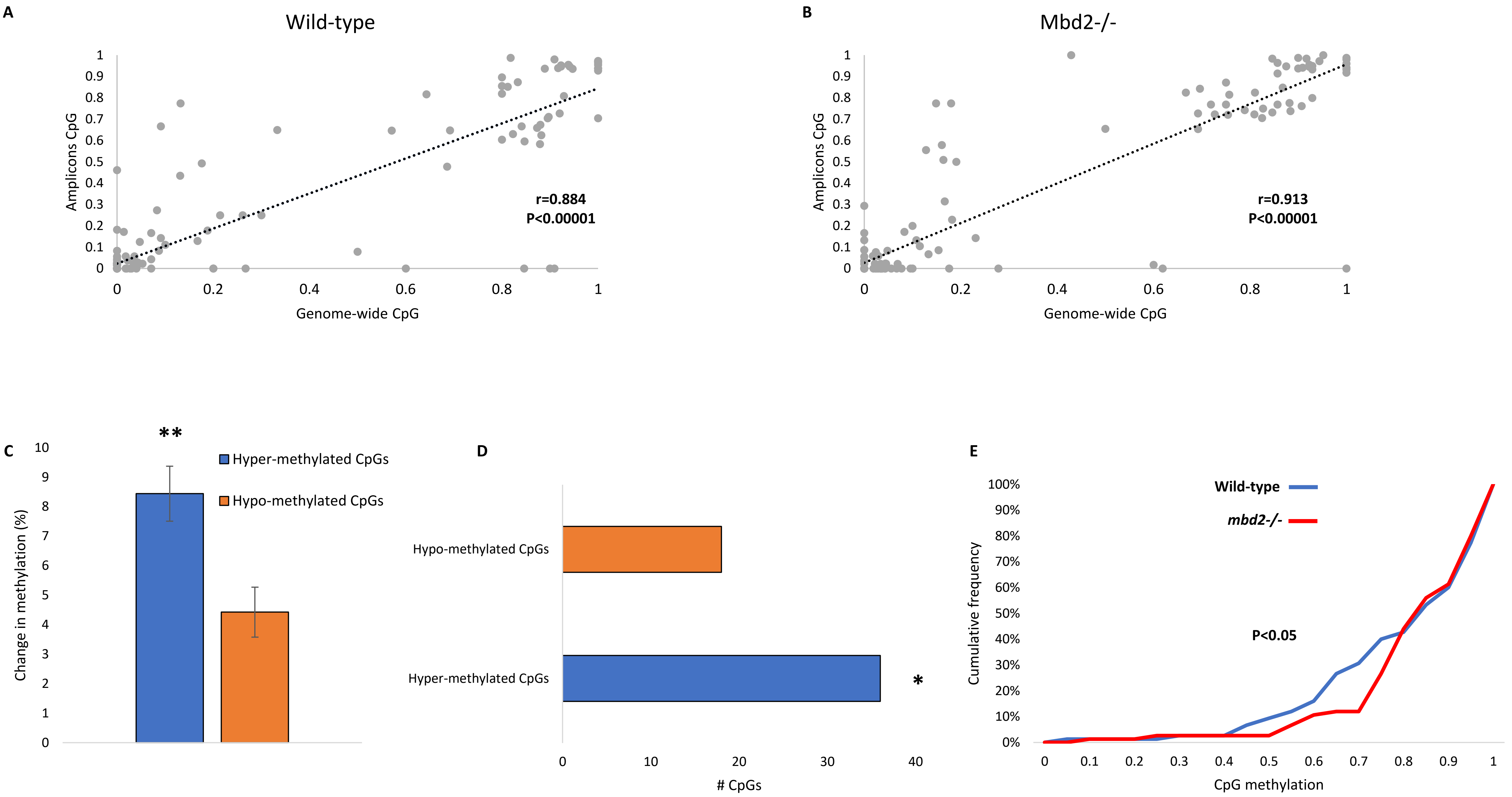

### Supplemental Figure 14

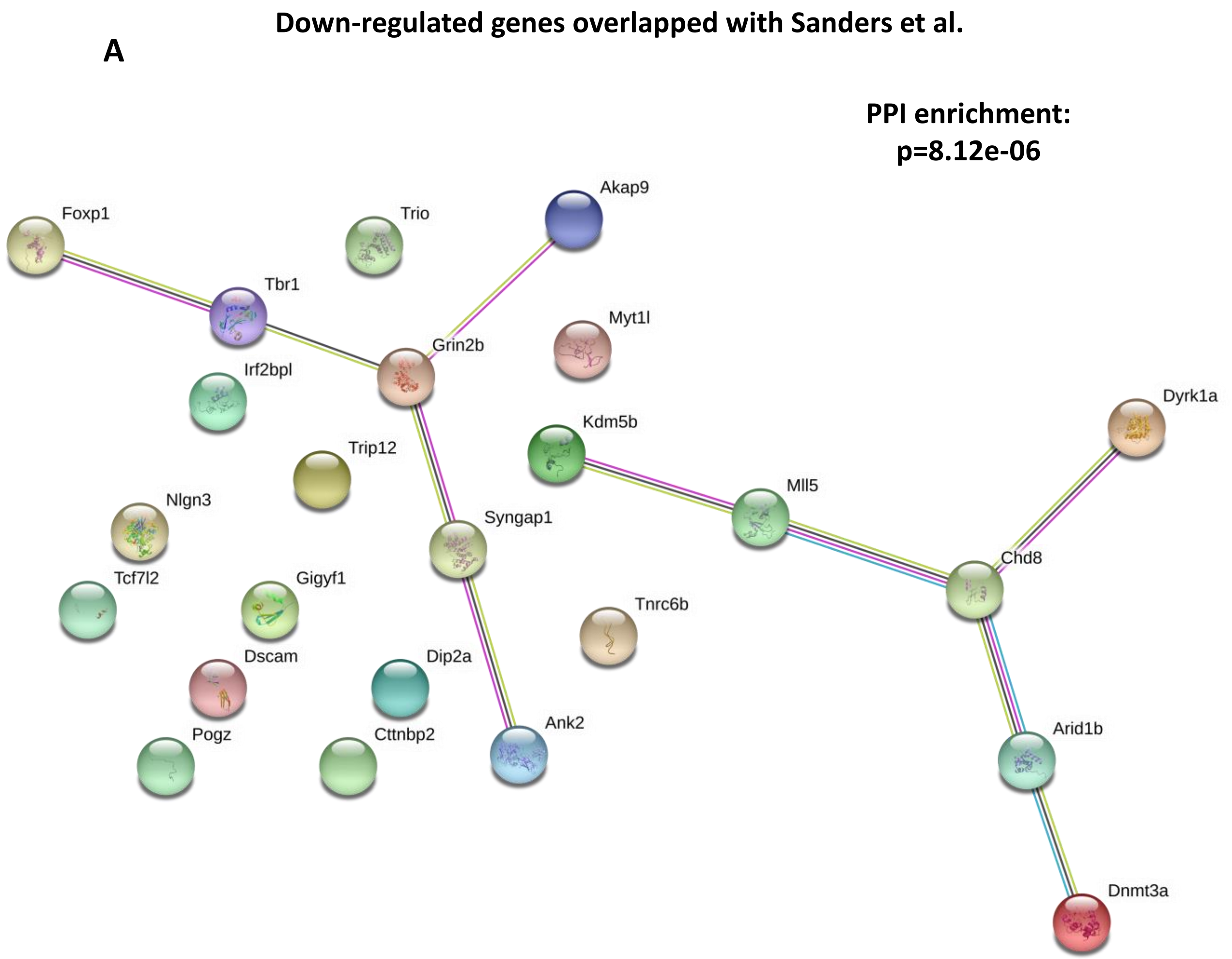

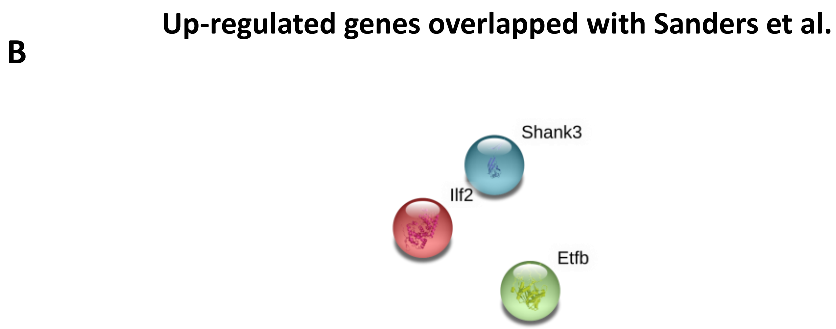

### Supplemental Figure 16

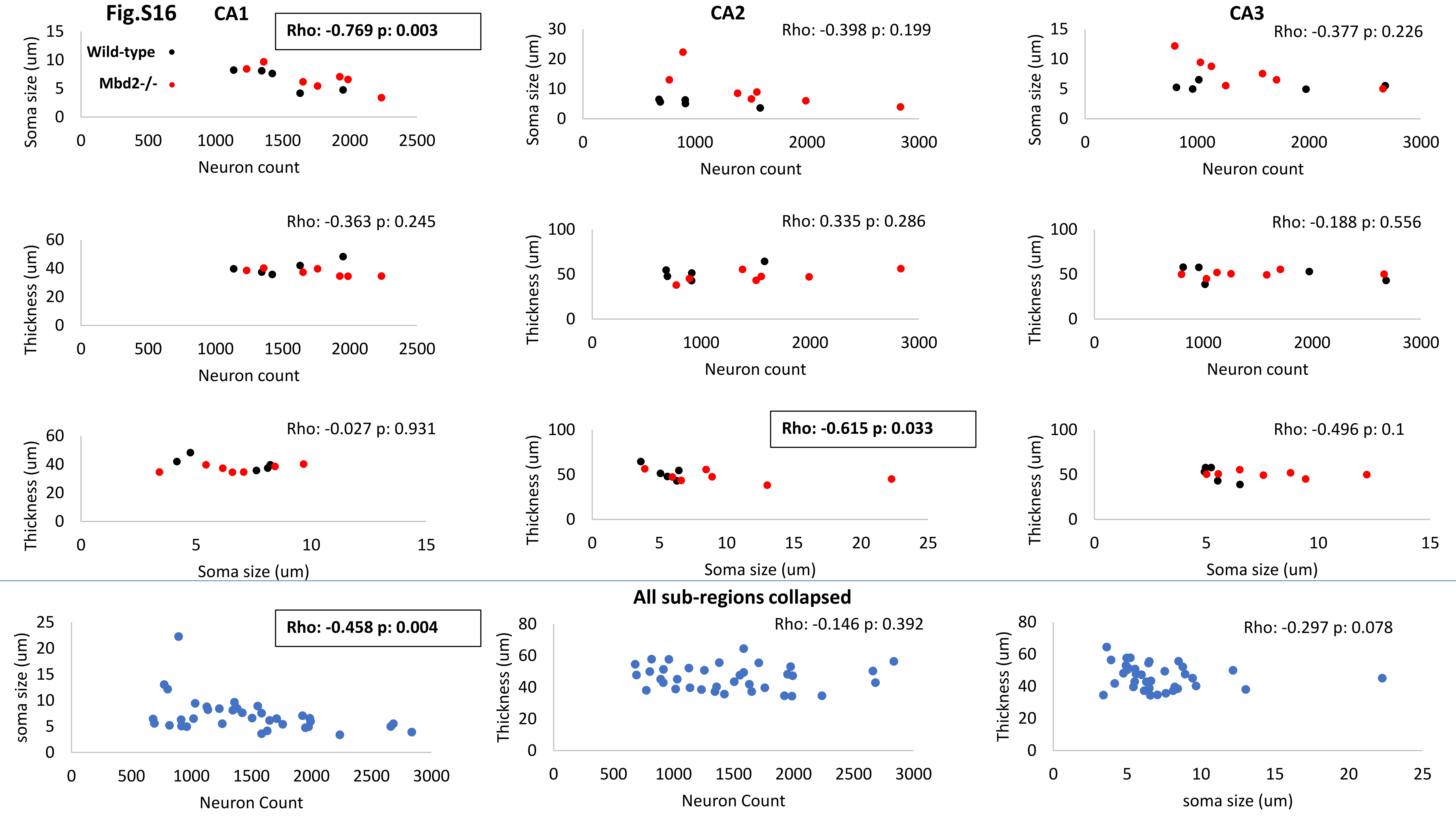

### Supplemental Figure 17

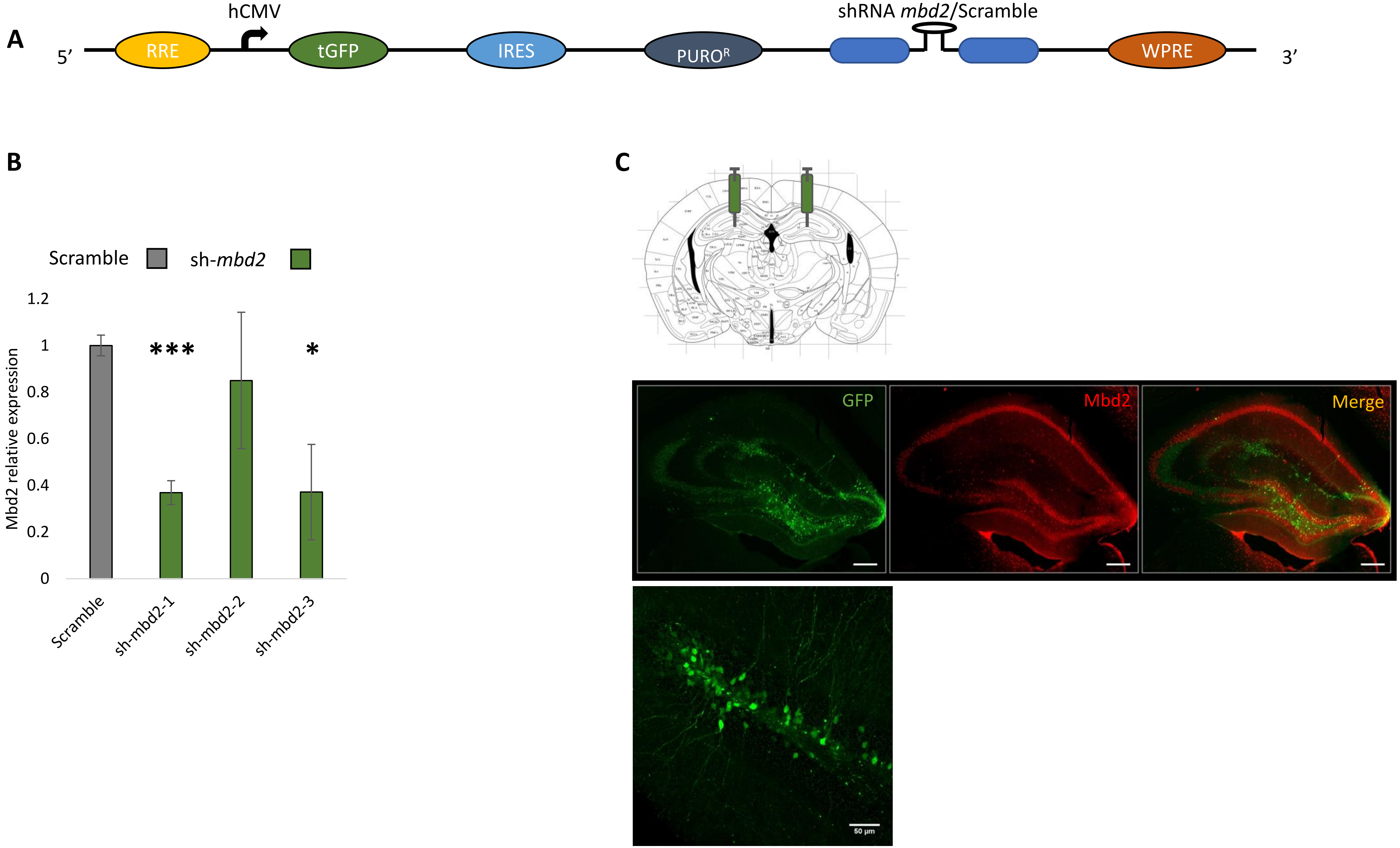
