## Supplemental Figure 5 for "Methyl-CpG binding domain 2 (Mbd2) is an Epigenetic Regulator of Autism-Risk Genes and Cognition"

**Fig.S5**

**A**

**De-novo motif (top 6)**

| Rank | Motif | P-value | log P-pvalue | % of Targets | % of Background | STD(Bg STD) | Best Match/Details |
|---|---|---|---|---|---|---|---|
| 1 | 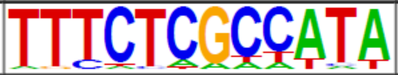 | 1e-149 | -3.450e+02 | 2.08% | 0.00% | 57.1bp (0.0bp) | E2F8/MA0865.1/Jaspar(0.748) More Information \| Similar Motifs Found |
| 2 | 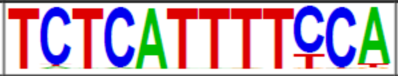 | 1e-108 | -2.492e+02 | 2.26% | 0.02% | 54.4bp (53.4bp) | NFAT5/MA0606.1/Jaspar(0.734) More Information \| Similar Motifs Found |
| 3 | 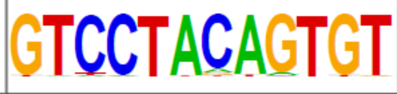 | 1e-104 | -2.399e+02 | 2.80% | 0.05% | 55.9bp (49.2bp) | ZKSCAN1(Zf)/HepG2-ZKSCAN1-ChIP-Seq(Encode)/Homer(0.656) More Information \| Similar Motifs Found |
| 4 | 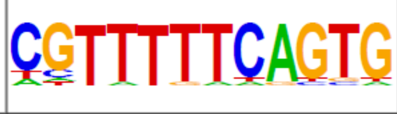 | 1e-81 | -1.887e+02 | 2.08% | 0.03% | 56.9bp (56.5bp) | Hoxd12(Homeobox)/ChickenMSG-Hoxd12.Flag-ChIP-Seq(GSE86088)/Homer(0.672) More Information \| Similar Motifs Found |
| 5 | 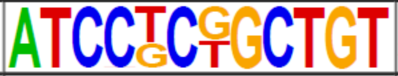 | 1e-35 | -8.164e+01 | 0.61% | 0.00% | 48.9bp (0.0bp) | Tcf12/MA0521.1/Jaspar(0.628) More Information \| Similar Motifs Found |
| 6 | 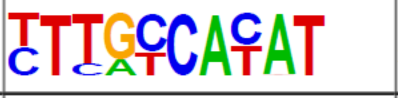 | 1e-33 | -7.810e+01 | 2.91% | 0.51% | 54.1bp (57.6bp) | CEBP:AP1(bZIP)/ThioMac-CEBPb-ChIP-Seq(GSE21512)/Homer(0.690) More Information \| Similar Motifs Found |

**B**

**Known motifs**

| Rank | Motif | Name | P-value | log P-pvalue | q-value (Benjamini) | # Target Sequences with Motif | % of Targets Sequences with Motif | # Background Sequences with Motif | % of Background Sequences with Motif |
|---|---|---|---|---|---|---|---|---|---|
| 1 | 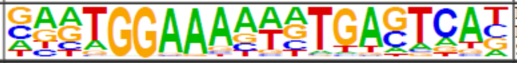 | NFAT:AP1(RHD,bZIP)/Jurkat-NFATC1-ChIP-Seq(Jolma_et_al.)/Homer | 1e-8 | -1.895e+01 | 0.0000 | 86.0 | 3.09% | 733.3 | 1.56% |
| 2 | 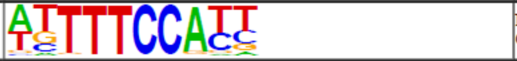 | NFAT(RHD)/Jurkat-NFATC1-ChIP-Seq(Jolma_et_al.)/Homer | 1e-5 | -1.341e+01 | 0.0003 | 337.0 | 12.11% | 4416.4 | 9.41% |
| 3 | 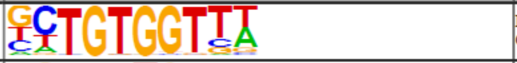 | RUNX-AML(Runt)/CD4+-PolII-ChIP-Seq(Barski_et_al.)/Homer | 1e-5 | -1.239e+01 | 0.0006 | 263.0 | 9.45% | 3361.1 | 7.16% |
| 4 | 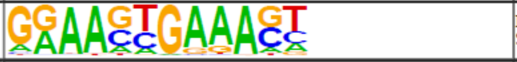 | IRF8(IRF)/BMDM-IRF8-ChIP-Seq(GSE77884)/Homer | 1e-3 | -8.070e+00 | 0.0335 | 114.0 | 4.10% | 1374.8 | 2.93% |
| 5 | 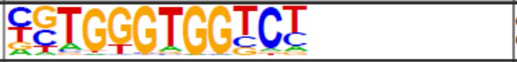 | Gli2(Zf)/GM2-Gli2-ChIP-Chip(GSE112702)/Homer | 1e-3 | -7.895e+00 | 0.0335 | 88.0 | 3.16% | 1013.7 | 2.16% |
| 6 | 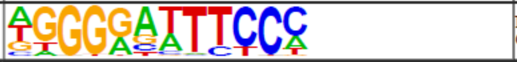 | NFkB-p65(RHD)/GM12787-p65-ChIP-Seq(GSE19485)/Homer | 1e-3 | -7.214e+00 | 0.0525 | 156.0 | 5.61% | 2025.5 | 4.32% |
| 7 | 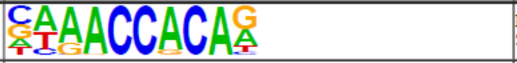 | RUNX(Runt)/HPC7-Runx1-ChIP-Seq(GSE22178)/Homer | 1e-2 | -5.769e+00 | 0.1909 | 244.0 | 8.77% | 3456.2 | 7.37% |
| 8 | 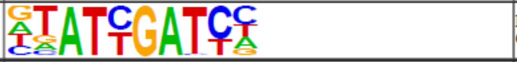 | HNF6(Homeobox)/Liver-Hnf6-ChIP-Seq(ERP000394)/Homer | 1e-2 | -5.241e+00 | 0.2832 | 178.0 | 6.40% | 2472.6 | 5.27% |
| 9 | 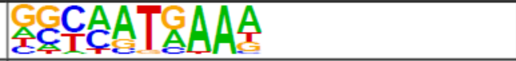 | Hoxd10(Homeobox)/ChickenMSG-Hoxd10.Flag-ChIP-Seq(GSE86088)/Homer | 1e-2 | -5.149e+00 | 0.2832 | 373.0 | 13.41% | 5546.9 | 11.82% |
| 10 | 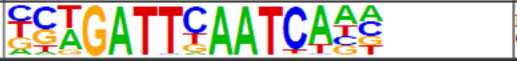 | DUX(Homeobox)/C2C12-Dux-ChIP-Seq(GSE87279)/Homer | 1e-2 | -5.103e+00 | 0.2832 | 4.0 | 0.14% | 12.4 | 0.03% |
| 11 | 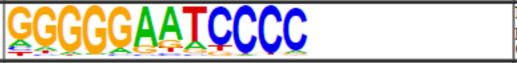 | NFkB-p50,p52(RHD)/Monocyte-p50-ChIP-Chip(Schreiber_et_al.)/Homer | 1e-2 | -5.093e+00 | 0.2832 | 31.0 | 1.11% | 318.6 | 0.68% |
| 12 | 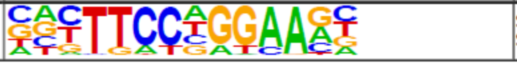 | Stat3+il21(Stat)/CD4-Stat3-ChIP-Seq(GSE19198)/Homer | 1e-2 | -4.874e+00 | 0.2832 | 218.0 | 7.84% | 3119.7 | 6.65% |
