## Supplemental Figure 12 for "Methyl-CpG binding domain 2 (Mbd2) is an Epigenetic Regulator of Autism-Risk Genes and Cognition"

### Fig.S12

**A**

| | Hypo-methylated CpGs | Hyper-methylated CpGs |
|---|---|---|
| **All CpGs (FDR<0.05)** | 133 (103 genes) | 190 (119 genes) |
| **All CpGs (p<0.001)** | 1519 (1330 genes) | 1486 (1214 genes) |
| **promoters (p<0.001)** | 494 (460 genes) | 476 (434 genes) |

**B**

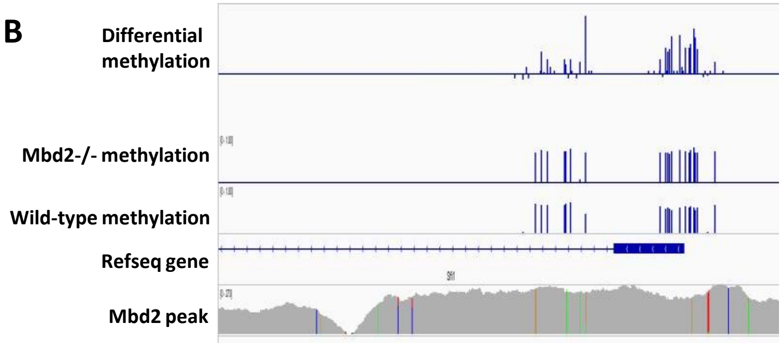


**C**

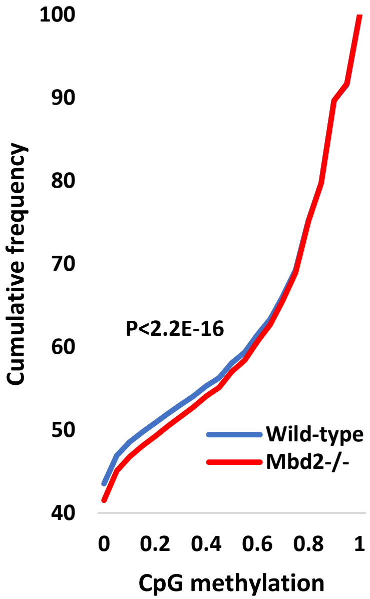


**D**

| | | | |
|---|---|---|---|
| oligos_6nt_mkv1_m1 | K-mer sig. 1.74<br>**e-value: 0.018** | versus jaspar_core_nonredundant_vertebrates:<br>**SP8, KLF14, SP3** | 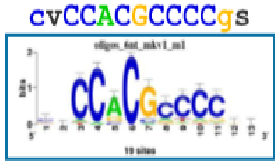 |
| oligos_7nt_mkv1_m1 | K-mer sig. 1.81<br>**e-value: 0.015** | versus jaspar_core_nonredundant_vertebrates:<br>**KLF14, Klf12, SP8** | 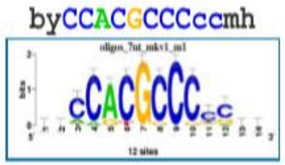 |
| oligos_7nt_mkv1_m2 | K-mer sig. 1.60<br>**e-value: 0.025** | versus jaspar_core_nonredundant_vertebrates: no match | 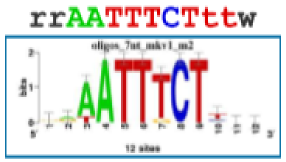 |

**E** **Hypo-methylated promoters (460 genes)**

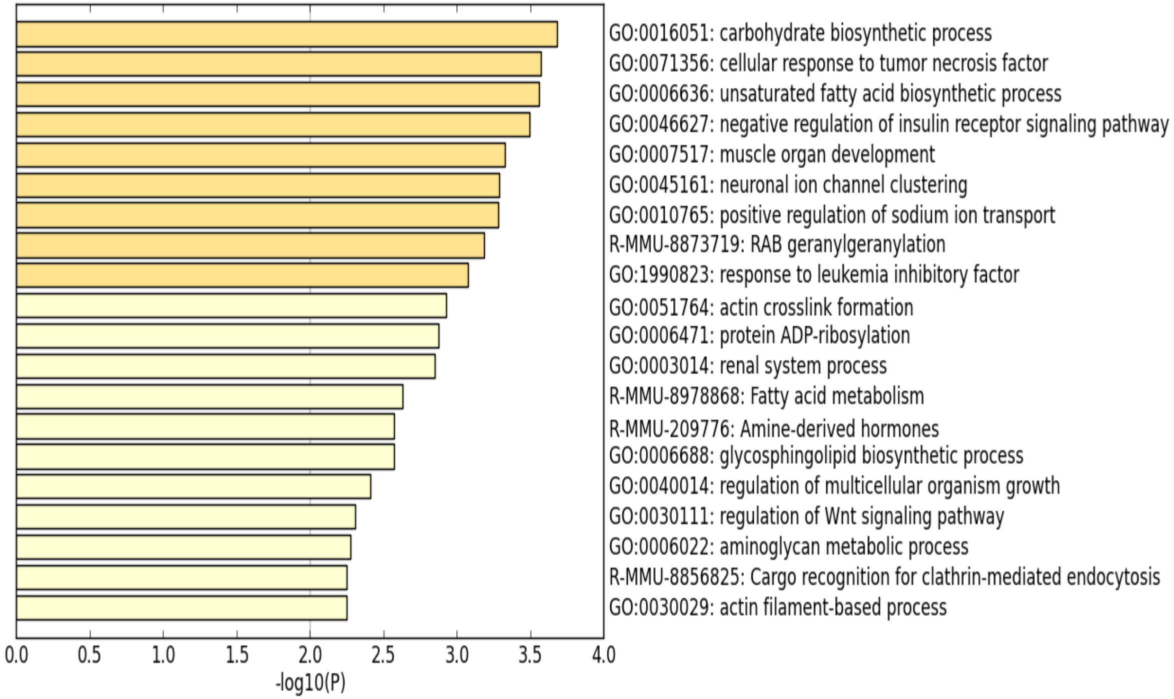


**F** **Hyper-methylated promoters (434 genes)**

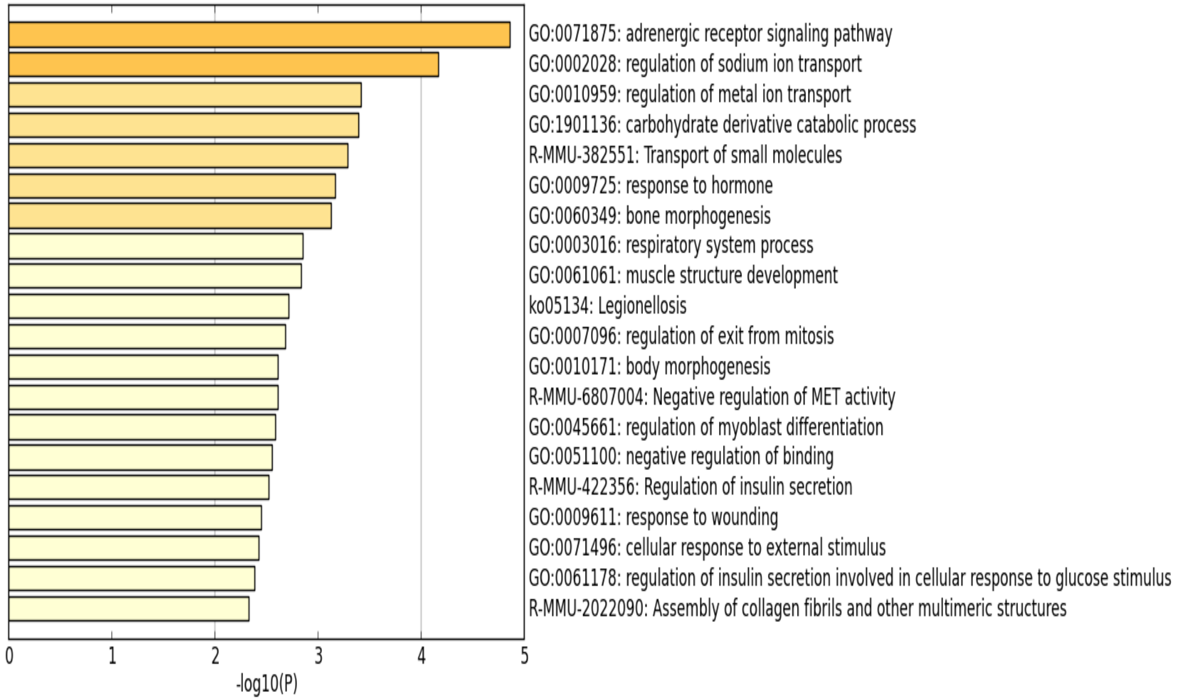
