## Supplemental Figure 15 for "Methyl-CpG binding domain 2 (Mbd2) is an Epigenetic Regulator of Autism-Risk Genes and Cognition"

Fig.S15

A

Down-regulated genes overlapped with SFARI-GENE dataset

PPI enrichment: p=1E-16

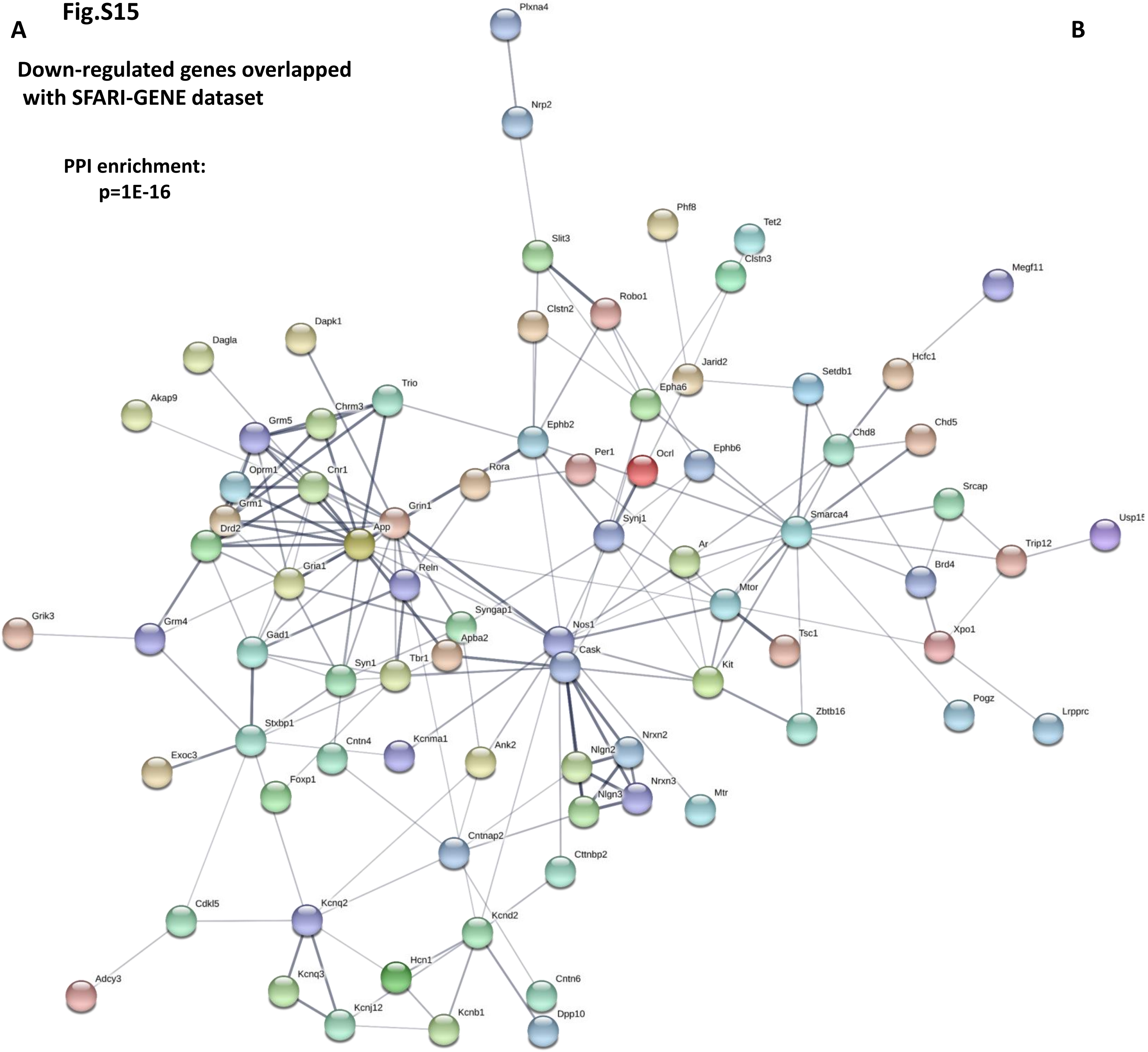


B

Up-regulated genes overlapped with SFARI-GENE dataset

PPI enrichment: p=0.00559

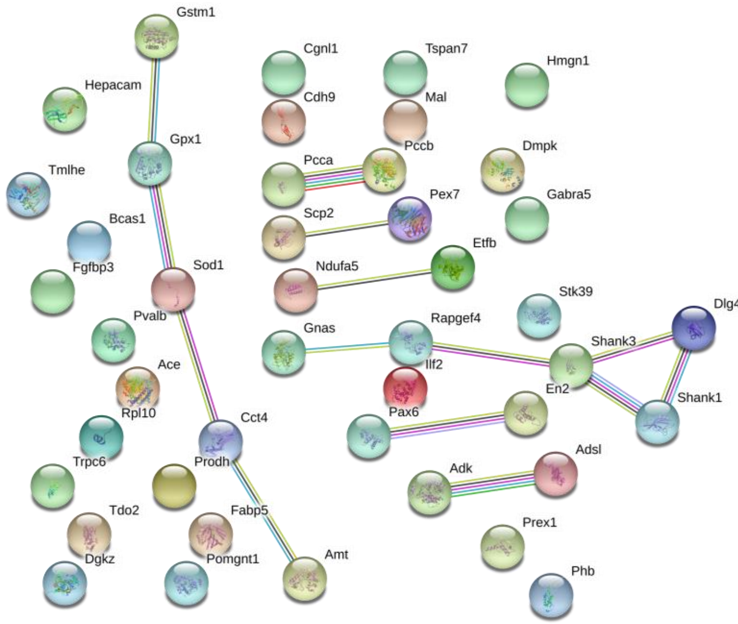
